# Presynaptic accumulation of α-synuclein causes synaptopathy and progressive neurodegeneration

**DOI:** 10.1101/2020.10.12.335778

**Authors:** Jessika C. Bridi, Erika Bereczki, Saffron K. Smith, Gonçalo M. Poças, Benjamin Kottler, Pedro M. Domingos, Christopher J. Elliott, Dag Aarsland, Frank Hirth

**Author notes:** Laboratory of Molecular and Functional Neurobiology, Department of Pharmacology, Institute of Biomedical Sciences, University of São Paulo, São Paulo, Brazil (Jessika C. Bridi). Correspondence to: Dr. Jessika Bridi;, Dr. Frank Hirth.

## Abstract

Alpha-synuclein (α-syn) mislocalisation and accumulation in intracellular inclusions is the major pathological hallmark of degenerative synucleinopathies, including Parkinson’s disease, Parkinson’s disease with Dementia and Dementia with Lewy Bodies. Typical symptoms are behavioural abnormalities including motor deficits that mark disease progression, while non-motor symptoms and synaptic deficits are already apparent during the early stages of disease. Synucleinopathies have therefore been considered synaptopathies that exhibit synaptic dysfunction prior to neurodegeneration. However, the mechanisms and events underlying synaptopathy are largely unknown. Here we investigated the cascade of pathological events underlying α-syn accumulation and toxicity in a *Drosophila* model of synucleinopathy by employing a combination of histological, biochemical, behavioural and electrophysiological assays. Our findings demonstrate that targeted expression of human α-syn leads to its accumulation in presynaptic terminals that caused downregulation of synaptic proteins, Cysteine String Protein, Synapsin, and Syntaxin 1A, and a reduction in the number of Bruchpilot puncta, the core component of the presynaptic active zone essential for its structural integrity and function. These α-syn-mediated presynaptic alterations resulted in impaired neuronal function, which triggered behavioural deficits in ageing *Drosophila* that occurred prior to progressive degeneration of dopaminergic neurons. Comparable alterations in presynaptic active zone protein were found in patient brain samples of Dementia with Lewy Bodies. Together, these findings demonstrate that presynaptic accumulation of α-syn impairs the active zone and neuronal function, which together cause synaptopathy that results in behavioural deficits and the progressive loss of dopaminergic neurons. This sequence of events resembles the cytological and behavioural phenotypes that characterise the onset and progression of synucleinopathies, suggesting that α-syn mediated synaptopathy is an initiating cause of age-related neurodegeneration.

## Introduction

Synucleinopathies are characterised by intraneuronal inclusions called Lewy bodies (LB) that are mainly formed of misfolded and aggregated forms of the presynaptic protein α-synuclein (α-syn) (Spillantini *et al.*, 1997; Bendor *et al.*, 2013; Walker *et al.*, 2015; Goedert *et al.*, 2017). These include Dementia with Lewy bodies (DLB) as well as and Parkinson’s disease with Dementia (PDD) and Parkinson’s disease (PD). PD is mainly characterised by the progressive loss of dopaminergic (DA) neurons in the substantia nigra pars compacta (SN), thereby depleting dopamine levels in synaptic terminals of the dorsal striatum (Dauer and Przedborski, 2003; Cheng *et al.*, 2010; Bridi and Hirth, 2018). The resulting regulatory imbalance in the basal ganglia causes a range of behavioural symptoms including bradykinesia, uncontrollable tremor at rest, postural impairment, and rigidity (Lang and Lozano, 1998*a*, *b*; Mochizuki *et al.*, 2018).

Although the majority of PD cases are sporadic, several genes including *LRRK2, Parkin, PINK1, DJ-1, GBA* and *SNCA* contribute to heritable cases of the disease (Klein and Westenberger, 2012; Ferreira and Massano, 2017). However, among the PD-related genes identified, the *SNCA* gene encoding α-syn remains the most potent culprit underlying PD, with a key pathogenic role both in familial and sporadic cases (Satake *et al.*, 2009; Simón-Sánchez *et al.*, 2009). Several point mutations in *SNCA* and increased gene dosage caused by duplication or triplication of the gene locus, are causally related to severe forms of PD (Singleton *et al.*, 2003; Kalia and Lang, 2015). These findings suggest a causal relationship between α-syn levels and the severity of cognitive decline, motor and non-motor symptoms, and neurodegeneration (Venda *et al.*, 2010; Bridi and Hirth, 2018).

Although the mechanisms underlying α-syn toxicity remain unclear, proteinaceous inclusions enriched with α-syn were found not only in LB within the neuronal soma but also in axonal processes (Braak *et al.*, 1999). Most importantly, α-syn micro-aggregates were found to be enriched in the presynaptic terminals of DLB patients (Kramer and Schulz-Schaeffer, 2007) along with phosphorylated α-syn, which is believed to disrupt synaptic structure and function (Colom-Cadena *et al.*, 2017). These findings suggest that α-syn accumulation may cause a toxic gain of function phenotype at the synapse, which impairs its function and connectivity, ultimately causing synaptopathy.

In line with this hypothesis, classical motor symptoms in PD become clinically apparent only when 60% of DA striatal terminals were already lost, while the loss of DA neurons in the SN is only around 30% (Burke and O’Malley, 2013). Corroborating these observation, it is well acknowledged that the onset of PD initiates at least 20 years prior to the detection of classical motor phenotypes, a period known as the prodromal phase (Hawkes *et al.*, 2010; Kalia and Lang, 2015; Mahlknecht *et al.*, 2015). It has been suggested that during this phase, a large number of proteins involved in synaptic transmission are affected, as indicated by their altered expression levels in PD and DLB patients (Bereczki et al., 2016; Dijkstra et al., 2015). These findings are in agreement with positron emission tomography of early-stage PD patients who presented extensive axonal damage and diminished nigrostriatal pathway connectivity (Caminiti *et al.*, 2017). Therefore, it has been hypothesised that neurodegeneration in PD and DLB follows a dying back-like pathomechanism, where degeneration of synapses and axonal connections precedes the loss of neurons, classifying them as synaptopathies (Calo *et al.*, 2016; Bridi and Hirth, 2018). However, it remains unclear how α-syn accumulation impairs synaptic homeostasis, its structure and function, ultimately leading to neurodegeneration.

Here we investigated the succession of events caused by cell and tissue-specific accumulation of α-syn. We employed a *Drosophila* model of synucleinopathy that expresses human wild type α-syn, and analysed post-mortem tissue of PD and DLB patients. Our findings demonstrate that α-syn accumulates in, and alters the presynaptic terminal, especially the active zone, which was also observed in the prefrontal cortex of DLB patients. In *Drosophila*, these alterations caused neuronal dysfunction and behavioural deficits that preceded degenerative loss of DA neurons - cytological and behavioural phenotypes that resemble the onset and progression of synucleinopathies. Together these findings provide experimental evidence that presynaptic accumulation of α-syn causes synaptopathy and progressive neurodegeneration.

## Results

### α-syn accumulates in presynaptic terminals

To investigate the consequences of α-syn accumulation on synapse structure and function, we used transgenic *Drosophila* to express human wild type α-syn fused to EGFP (*UAS-WT-α-syn-EGFP*) alongside with control animals expressing EGFP only (*UAS-EGFP*) (Poças *et al.*, 2015). This genetic model is an invaluable tool to investigate the toxic gain-of-function of α-syn as the *Drosophila* genome lacks a homologue of the *SNCA* gene (Feany and Bender, 2000). We first compared the expression pattern of α-syn to the respective EGFP control by using the pan-neuronal driver *nSyb-Gal4.* Using immunohistochemistry, the localisation and expression pattern of α-syn were determined at the *Drosophila* L3 larval neuromuscular junction (NMJ) by colocalisation with Cysteine String Protein (CSP) and horseradish peroxidase (HRP) (Fig. 1A-B). CSP labels boutons containing synaptic vesicles at the presynaptic terminal (Fernández-Chacón *et al.*, 2004; Burgoyne and Morgan, 2015) while HRP labels synaptic membranes (Menon *et al.*, 2013) (Fig. 1A). The control expressing EGFP only and the experimental group expressing α-syn-EGFP displayed co-localisation with CSP (Fig. 1B). In addition, EGFP and α-syn were also detectable in axonal domains labelled with anti-HRP and devoid of synaptic boutons, however, the EGFP expression pattern in these areas was very distinct between control and experimental group (Fig. 1B-C, arrowheads and dashed boxes). α-syn immunolabelling was enriched in regions with a high density of synaptic boutons. In order to quantify this phenotype, EGFP intensity was measured in regions of CSP-positive synaptic boutons, and in CSP-negative axonal regions, in both control and experimental conditions. A ratio of EGFP intensity was calculated by dividing the values obtained from synaptic boutons by values obtained from axonal regions. Values around 1 indicated that EGFP intensity is similar in the two regions while values higher than 1 identified a higher EGFP intensity in synaptic boutons compared to axonal regions. The analysis revealed that α-syn immunofluorescence was significantly higher in synaptic boutons than in axons compared to the control NMJ expressing EGFP only (Fig. 1B-C and Table S1, arrowheads and dashed boxes). These data suggest that α-syn accumulates predominantly in presynaptic terminals and to a lesser extent in axonal areas devoid of synaptic boutons in the NMJ.

**Figure 1.**
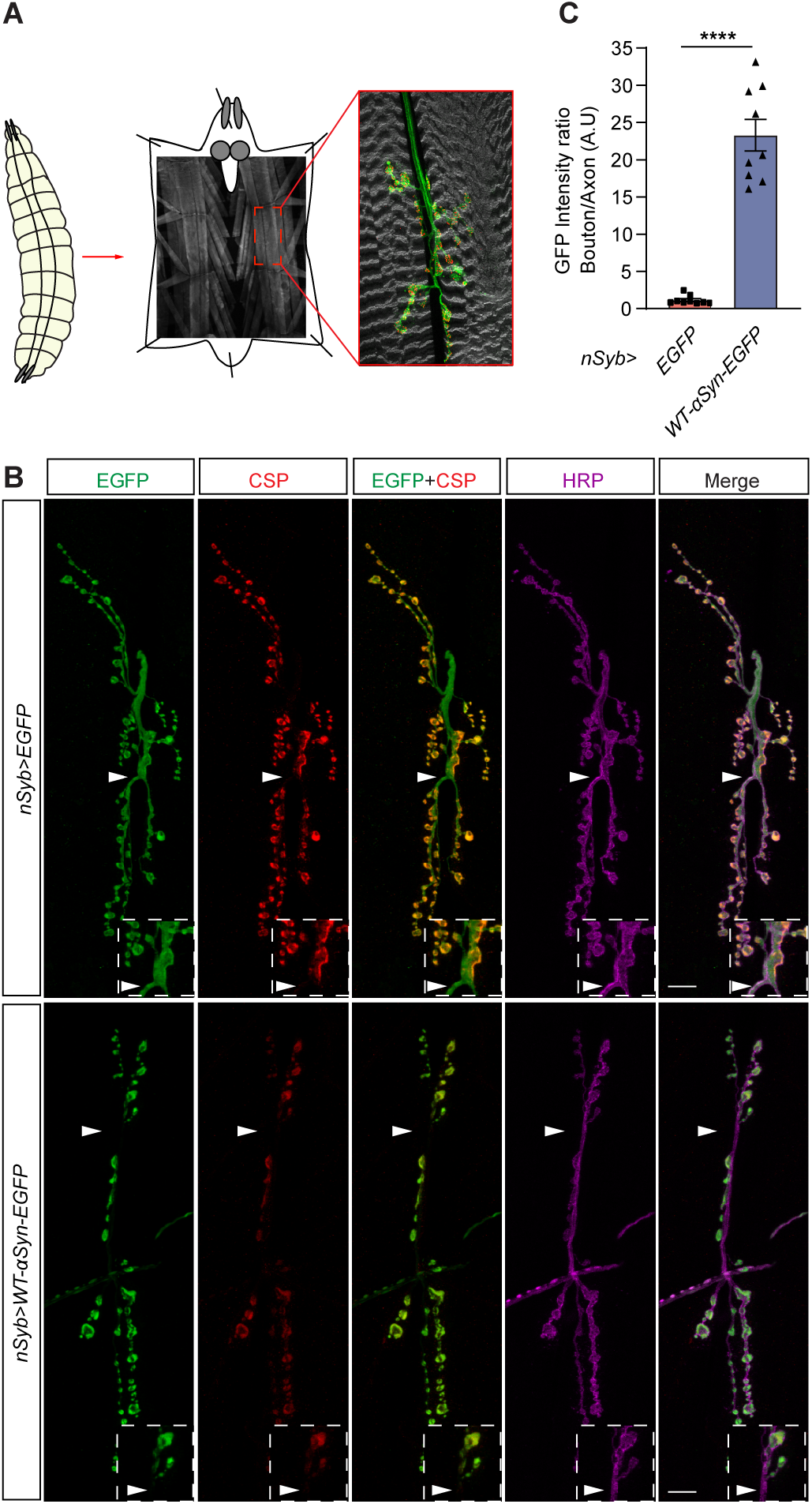
α-syn accumulates in synaptic boutons at the *Drosophila* NMJ. (A) Third instar larval stage (L3) *Drosophila* (left) was used to investigate presynaptic terminals (boutons) terminating at the neuromuscular junction (NMJ, middle). Immunohistochemistry assay reveals the muscle fibres (grey) stained with phalloidin that binds to actin; the axons descending from the motor neuron and terminating onto the muscles are labelled with anti-horseradish peroxidase (HRP – grey); and the presynaptic terminal of the motor neuron labelled with anti-BRP (red). (B) Representative image of NMJ of *nSyb*>*EGFP* and *nSyb*>*WT-α-syn-EGFP* larvae immunostained with anti-GFP (green), anti-CSP (red) and anti-HRP (magenta). Arrowheads indicate accumulation of WT-α-syn-EGFP in synaptic boutons, which are immunolabelled with anti-CSP while control EGFP is homogeneously expressed in synaptic boutons and axonal regions devoid of CSP immunoreactivity; dashed boxes show a higher magnification of the areas indicated by the arrowheads (C) Quantitative analysis of the ratio of GFP fluorescence intensity between boutons and axons; ****p= 0.0001; mean ± SEM shown for each genotype (n = 9 NMJs/genotype). Statistical analyses were performed using unpaired two-tailed t-test. Scale bars: 10 μm.

### Accumulation of α-syn affects presynaptic proteins

Previous studies demonstrated that the levels of synaptic proteins are altered in patients with PD and related disorders (Dijkstra *et al.*, 2015; Bereczki *et al.*, 2016), however, it is unclear whether and how α-syn accumulation might be related to the disease. We, therefore, investigated whether presynaptic accumulation of α-syn leads to alterations in the expression and/or localisation of presynaptic proteins that are essential for neurotransmission. We first used the larval L3 NMJ to evaluate the effect of α-syn accumulation on synaptic vesicle proteins CSP and Synapsin, the SNARE complex proteins, SNAP-25 and Synaptobrevin, and the synaptic vesicle-specific Ca^2+^ binding protein Synaptotagmin.

CSP is a synaptic vesicle protein whose loss-of-function in *Drosophila* causes synaptic degeneration and lethality (Zinsmaier *et al.*, 1990, 1994). CSP acts as a chaperone in the presynaptic terminal (Burré *et al.*, 2010; Sharma *et al.*, 2011) and has been shown to be altered in PD patients as well as in cellular models inoculated with pre-formed fibrils of α-syn (Volpicelli-Daley *et al.*, 2011; Dijkstra *et al.*, 2015). We measured the expression levels and localisation of CSP at the larval NMJ by immunofluorescence, which revealed that CSP levels but not its localisation were downregulated in synaptic boutons co-labelled with α-syn when compared to EGFP controls (Fig. 2A). Quantification of mean fluorescence intensity was performed as a proxy for expression level in single synaptic boutons revealed significantly reduced CSP levels in larvae expressing α-Syn compared to controls expressing EGFP only (Fig. 2C and Table S1).

**Figure 2.**
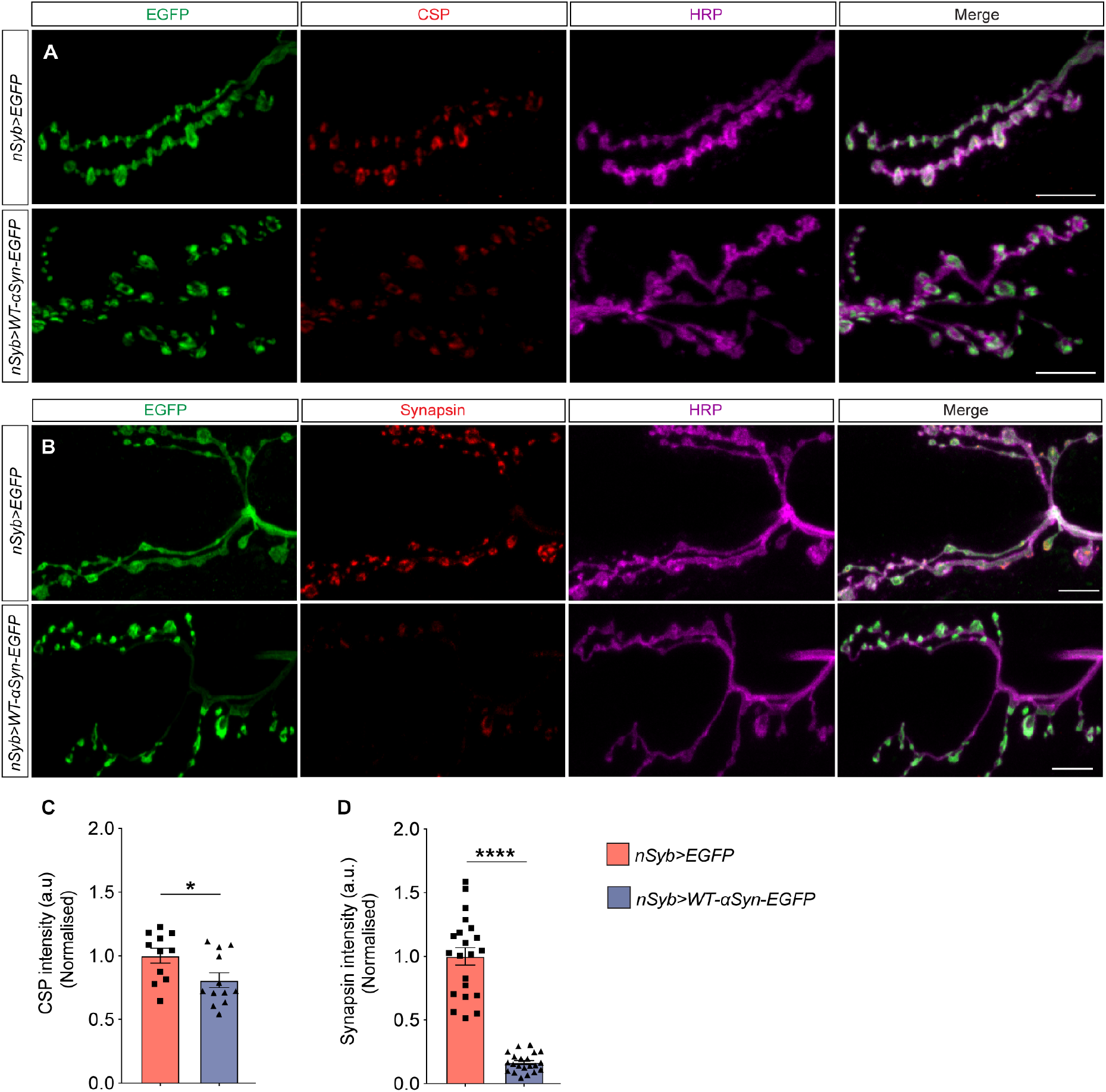
Accumulating α-syn downregulates presynaptic proteins. (A) Confocal images of NMJ immunolabeled with anti-CSP, anti-GFP and anti-HRP. (B) Confocal images of NMJ staining with anti-Synapsin, anti-GFP and anti-HRP. (C) Quantitative analysis of the fluorescence levels of CSP revealed downregulation in the NMJ of *nSyb*>*WT-α-syn-EGFP* larvae compared to control; *p= 0.0297; mean ± SEM shown for each genotype (n= 11 NMJs/ genotype). (D) Quantitative analysis of the Synapsin fluorescence levels showed downregulation in the NMJ of *nSyb*>*WT-α-syn-EGFP* larvae compared to control; ****p<0.0001; mean with SEM shown for each genotype (n= 21 NMJs/genotype). Statistical analyses were performed using unpaired two-tailed t-test. t Scale bars: 10 μm

We next examined the presynaptic protein Synapsin which is associated with the cytoplasmic surface of the synaptic vesicle membrane and plays a fundamental role in regulating vesicle trafficking (Greengard *et al.*, 1993; Gitler *et al.*, 2004; Südhof, 2004). We measured Synapsin fluorescence intensity and localisation in single synaptic boutons of larvae expressing either EGFP as control or α-syn (Fig. 2B). Quantitative analysis of Synapsin fluorescence intensity revealed that α-syn accumulation caused a reduction in Synapsin expression levels in synaptic boutons, compared to the control group (Fig. 2D and Table S1); however, alterations in Synapsin localisation were not observed. In contrast to CSP and Synapsin, we did not observe any changes in fluorescence intensity levels and localisation of SNAP-25, Synaptobrevin or Synaptotagmin (Fig. S1).

To examine the progression of α-syn-mediated presynaptic deficits, we assessed α-syn accumulation in adult flies at day 3 and 20 and performed western blots of heads of flies expressing α-syn or EGFP. While the levels of EGFP were unchanged (p=0.5777; Fig. 3A and Table S1), elevated levels of α-syn were increased over time (*p=0.0411; Fig. 3A and Table S1). We then assessed the levels of Synapsin and Syntaxin in adult flies, which revealed downregulated Synapsin levels at day 3 and day 20 (3-DO: *p=0.0328; 20-DO: **p=0.002; Fig. 3B and Table S1), similar to the phenotype observed in synaptic boutons of the NMJ. The expression levels of Syntaxin, a component of the SNARE complex (Han *et al.*, 2017), were reduced at day 3 but only significantly downregulated at day 20 when compared to control (3-DO: p=0.1776; 20-DO: **p=0.0093; Fig. 3C and Table S1). Together, these findings indicate that α-syn accumulates in the *Drosophila* brain, which in turn causes specific alterations of presynaptic proteins, with CSP, Syntaxin and Synapsin being especially susceptible to the deleterious effects of accumulating α-syn.

**Figure 3.**
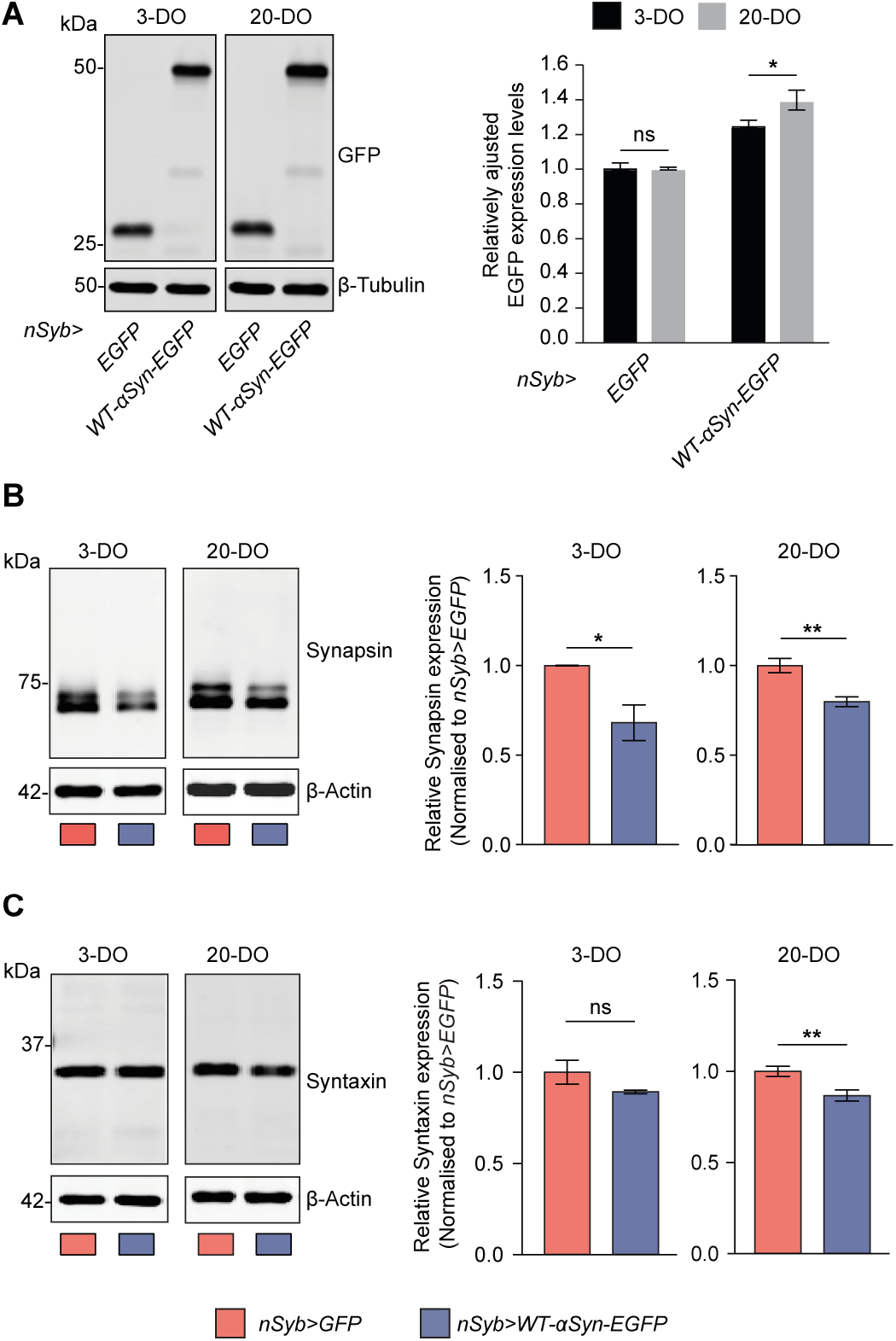
Progressive accumulation of α-syn affects presynaptic proteins in ageing animals. (A) Expression levels of WT-α-syn-EGFP expressed under the control of pan-neuronal driver *nSyb-Gal4* increased at 20 days of age compared to the levels observed in fly brains at 3 days of age; *p=0.0411, ns – not significant p>0.05, n=7-9. Such accumulation was specific for WT-α-syn-EGFP since the control flies expressing EGFP only showed no alteration in the expression levels of EGFP. (B) The expression levels of the synaptic vesicle protein Synapsin were reduced in flies expressing WT-α-syn-EGFP at day 3 and 20 compared to control flies; *p= 0.03 and **p= 0.002, n= 3-6. (C) The expression levels of Syntaxin, a protein of the presynaptic SNARE complex, also had its levels reduced at day 20 in fly brain expressing WT-α-syn-EGFP compared to control; **p= 0.0093, ns – not significant p>0.05, n=3-6. Mean ± SEM are shown, statistical analyses were performed using unpaired two-tailed t-test.

### Presynaptic accumulation of α-syn reduces active zone density

Given the impact of α-syn on presynaptic proteins, we investigated whether the accumulation of α-syn may alter the active zone (AZ) in *Drosophila.* The AZ is a specialised presynaptic site required for vesicle docking and neurotransmitter exocytosis, conveying speed and accuracy to synaptic transmission (Zhai and Bellen, 2004; Südhof, 2012; Wang *et al.*, 2016). We used confocal and instant structured illumination microscopy (iSIM) to NMJs labelled with anti-Bruchpilot (BRP) (Fig. 4A-B). BRP encodes a cytoskeletal protein essential for the structural integrity and the function of electron-dense projection (T-bar) at the AZ (Kittel *et al.*, 2006; Wagh *et al.*, 2006).

**Figure 4.**
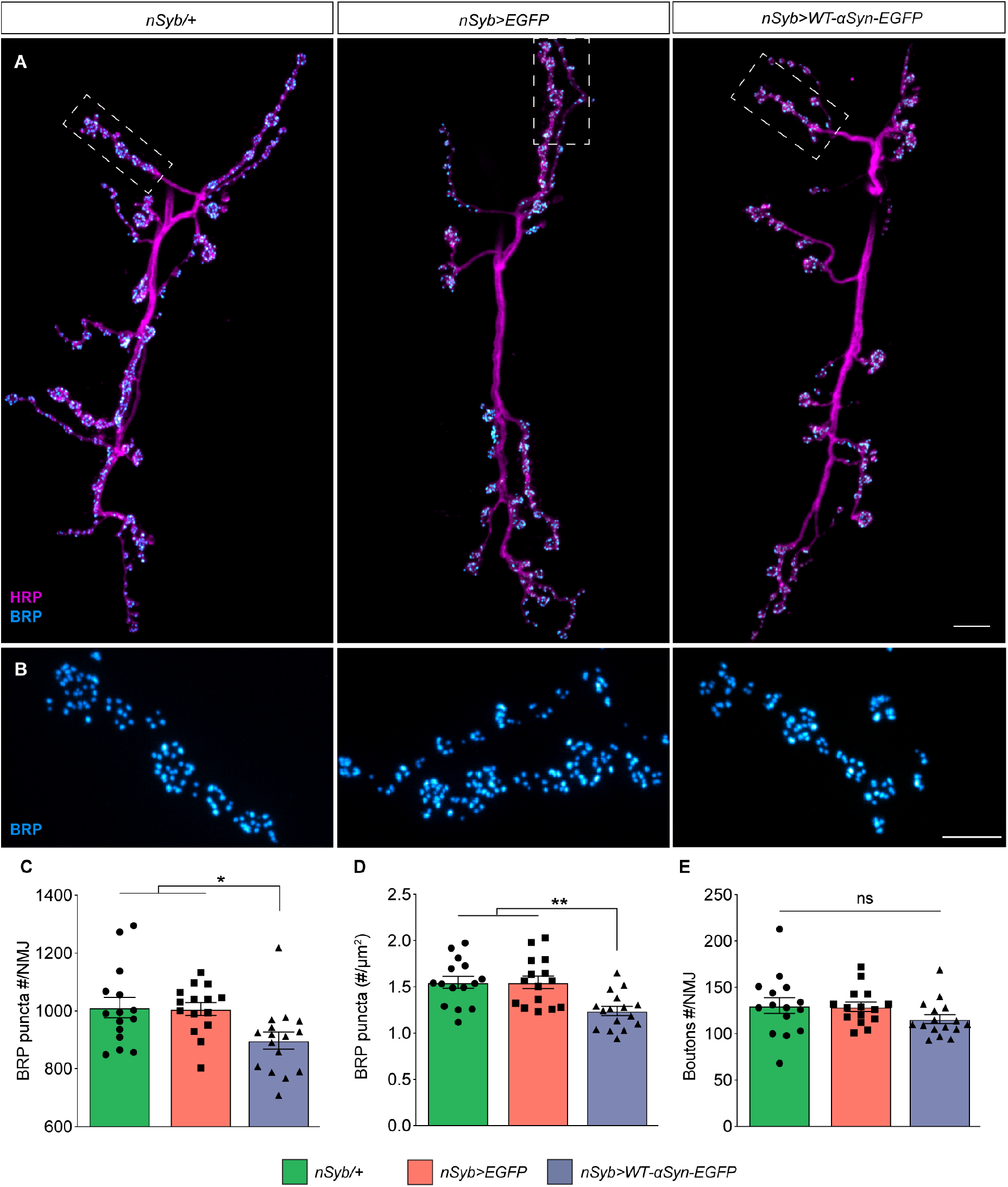
Presynaptic active zones are reduced by accumulating α-syn. (A) Confocal images of NMJ, immunolabeled with anti-nc82/BRP and anti-HRP. (B) Dashed box represents the section of the NMJ (A) imaged with instant super resolution structured illumination microscopy (iSIM) (C) Quantitative analysis of total number of BRP puncta that represents the AZs labelled with anti-nc82/BRP per NMJ; *p= 0.0245 compared to *nSyb*/+ and *p= 0.0317 compared to *nSyb*>*EGFP.* (D) Number of BRP puncta normalised by the synaptic surface μm^2^ immunolabelled with anti-HRP; **p= 0.0023. (E) The total number of synaptic boutons/NMJ was unchanged. All graphs are represented as mean ± SEM shown for each genotype. (n=15-16 NMJs/genotype). Statistical analyses were performed using one-way ANOVA test followed by Tukey’s multiple comparison post-hoc test. Scale bars: (A) 10 μm and (B) 5 μm.

Expression and accumulation of α-syn in *nSyb*>*WT-α-syn-EGFP* L3 larvae caused a reduction in the total number of BRP-labelled AZ puncta compared to control groups *nSyb*/+ and *nSyb*>*EGFP* (*vs. nSyb/+ *p=0.0245; vs. nSyb>EGFP *p=0.0317*; Fig. 4C and Table S1). A detailed analysis accounting also for the surface area of the NMJ labelled by anti-HRP (Goel and Dickman, 2018), revealed a more significant reduction in the total number of puncta in *nSyb*>*WT-α-syn-EGFP* larvae, when compared to controls (*vs. nSyb/+ and nSyb>EGFP **p=0.0023*; Fig. 4D and Table S1). In contrast, neither the total number of synaptic boutons (*vs. nSyb>EGFP p=0.3037*; Fig. 4E and Table S1) nor the morphology and localisation of BRP puncta (Fig. 4B – dashed lines) were affected. Together these data demonstrate that accumulation of α-syn alters the AZ of presynaptic terminals.

### α-syn accumulation impairs neuronal function

The core function of BRP is the maintenance of the presynaptic AZ and thus synaptic homeostasis and function (Wagh *et al.*, 2006). The observed reduction of BRP-positive puncta suggested that physiological defects could occur as a result of presynaptic accumulation of α-syn. To investigate whether synaptic efficacy and neurotransmission were affected, we determined the Steady State Visual Evoked Potential (SSVEP) in adult flies (Fig. 5). The SSVEP quantifies the physiological response to flickering stimuli which generate frequency- and phase-locked response components with a very high signal-to-noise ratio (Fig. 5A-C) (Afsari *et al.*, 2014; Petridi *et al.*, 2020). In *Drosophila*, the net response of flickering stimuli is mediated by retinal photoreceptor cells together with lamina and medulla neurons that are electrically linked (Fig. 5C-D) (Heisenberg, 1971). The visual response negatively correlates with dopamine levels in the brain (Calcagno *et al.*, 2013), with dopamine required to inhibit the response to visual stimuli (Afsari *et al.*, 2014).

**Figure 5.**
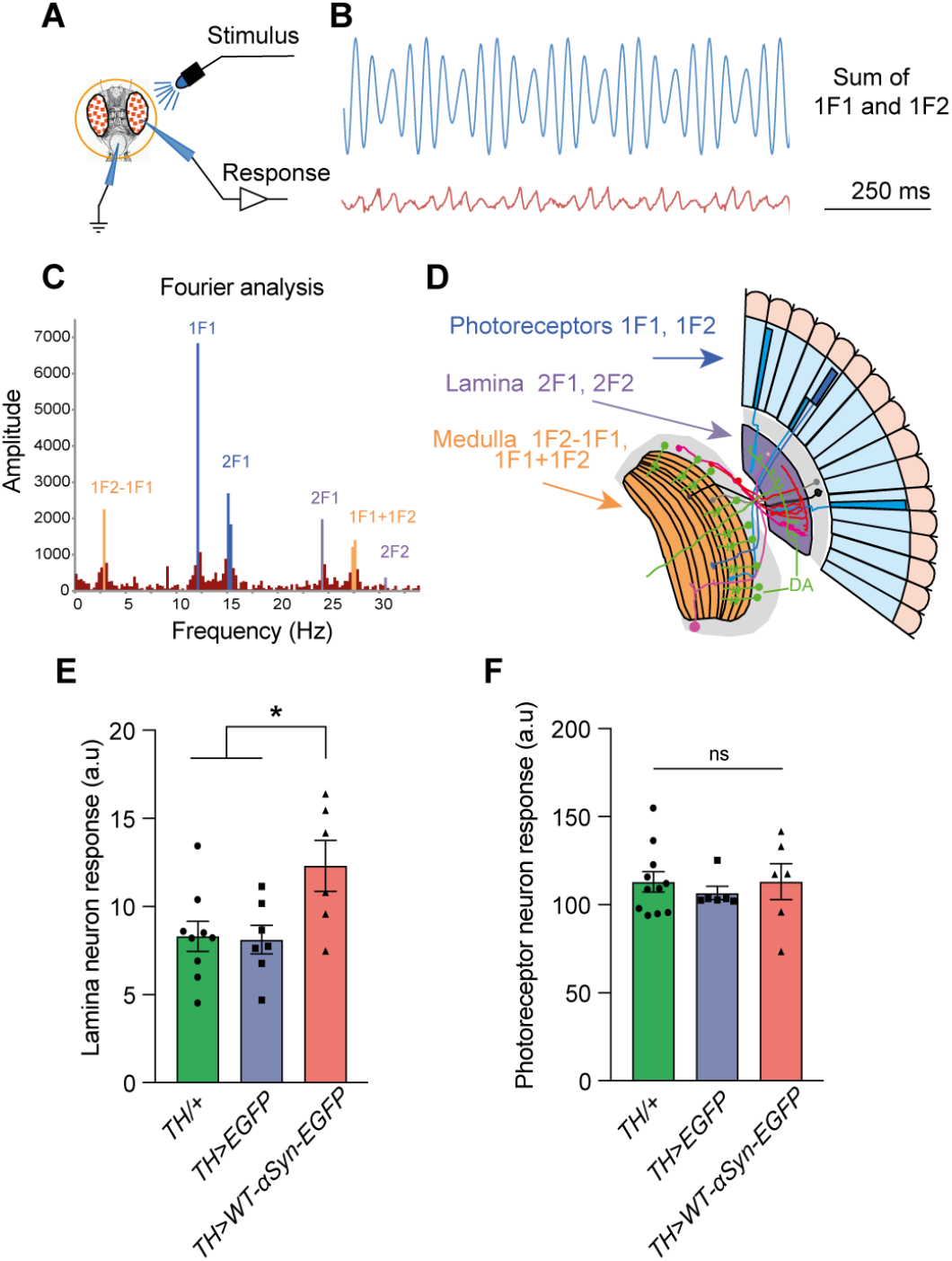
α-syn accumulation in dopaminergic neurons impairs visual response. (A) Steady-state visual evoked potential (SSVEP) was measured in flies raised in the dark and restrained in a Gilson pippete tip. The recording electrode was placed on the surface of the eye, with a reference electrode in the mouthparts. A full field blue light stimulus is provided from an LED. A pattern of 45 stimuli are provided, with different amounts of the 1F1 (12Hz) and 2F1 (15Hz) stimuli. (B) An example stimulus made up of 70% contrast at 1F1, and 30% contrast at 2F1 and its response. (C) The response (in B) is analysed by the FFT (Fast-Fourier Transform), revealing that the visual system responds to the supplied frequencies (1F1, 2F1) but also to their multiples (2F1, 2F2) and to the sums and differences (1F1+1F2; 1F2-1F1). Higher frequency harmonics are also seen, notably 2F1+2F2. (D) Genetic dissection (Afsari *et al.*, 2014; Nippe *et al.*, 2017) shows that the majority of the 1F1 and 1F2 components come from the photoreceptors, the 2F1 and 2F2 come from the lamina neurons, and the 1F1+1F2, 1F1+1F2 and 1F2-1F1 from the medulla. (A-D) Modified after Afsari *et al.*, 2014 and Petridi *et al.*, 2020. (E) Flies expressing WT-α-Syn-EGFP under control of *TH-Gal4* driver demonstrated a higher lamina neuron activity in the SSVEP, which is known to negatively correlate with levels of dopamine in the brain; *p= 0.03. (F) Photoreceptors of *TH*>*WT-α-Syn-EGFP* displayed no alteration in their response compared to control groups; ns – not significant p>0.05, n=6-11 flies/genotype. Mean ± SEM are shown, statistical analyses were performed using one-way ANOVA with Tukey’s multiple comparison post-hoc test.

Our analysis revealed that lamina neurons in 3-day-old *TH*>*WT-α-syn-EGFP* flies showed an increased SSVEP response upon stimulation when compared to control groups (*vs. TH*/+ *p=0.0313*; vs. TH>EGFP* *p=0.0325; Fig. 5E and Table S1). This response was specific to lamina neurons, as photoreceptor cells showed no alteration in their SSVEP response (*vs. TH*/+ p=0.9998*; vs. TH>EGFP* p=0.8261; Fig. 5F and Table S1). The difference in stimuli response between the lamina and photoreceptor neurons correlated well with the far more extensive DA innervation in the lamina than the photoreceptor layer (Fig. 5D) (Afsari *et al.*, 2014; Petridi *et al.*, 2020). Because dopamine inhibits the SSVEP response (Afsari *et al.*, 2014), these data suggest that *TH-Gal4* driven accumulation of α-syn in DA neurons impaired synaptic output of DA-rich lamina neurons, thereby reducing their visual response inhibition which in turn resulted in an increased SSVEP response.

### Accumulation of α-syn progressively impairs motor behaviour

PD patients suffer from a variety of motor symptoms ranging from resting tremor to bradykinesia and akinesia, related to the reduction of striatal dopamine (Mazzoni *et al.*, 2012). However, these symptoms only become clinically apparent when a large proportion of DA neurons have already been lost (Bridi and Hirth, 2018). We, therefore, investigated whether the accumulation of α-syn and in turn the alterations in presynaptic proteins would cause any motor deficits in adult flies. We employed three independent assays to quantify voluntary and reflexive motor behaviour, the *Drosophila* ARousal Tracking system (DART) (Faville *et al.*, 2015), startled-induced negative geotaxis (SING) (White *et al.*, 2010; Ruan *et al.*, 2015) and the proboscis extension response (PER) (Cording *et al.*, 2017).

We specifically targeted α-syn to DA neurons and measured their spontaneous motor activity with DART at 3 and 20 days of age (Fig. 6A). The activity and speed of *TH*>*WT-α-syn-EGFP* flies was greatly impaired at day 3 (Fig. 6B-C and Table S1) compared to control flies *TH-Gal4*/+ and *WT-α-syn-EGFP/+.* Notably, both activity and speed were further reduced in flies accumulating α-syn compared to controls at day 20 (Fig. 6B-C and Table S1). To better understand the detrimental impact of α-syn on movement patterns, we decomposed and analysed movement as units. The first unit comprised the initiation of a motor action (Fig. 6D, red arrows), the action initiation; the second unit was the bout length, depicting for how long the flies maintain a bout of activity (Fig. 6D, dark grey boxes); and the third unit was the inter-bout interval that quantifies the duration of the pause between the end of a previous and the beginning of a new bout of activity (Fig. 6D, white boxes). These units were collectively named activity metrics.

**Figure 6.**
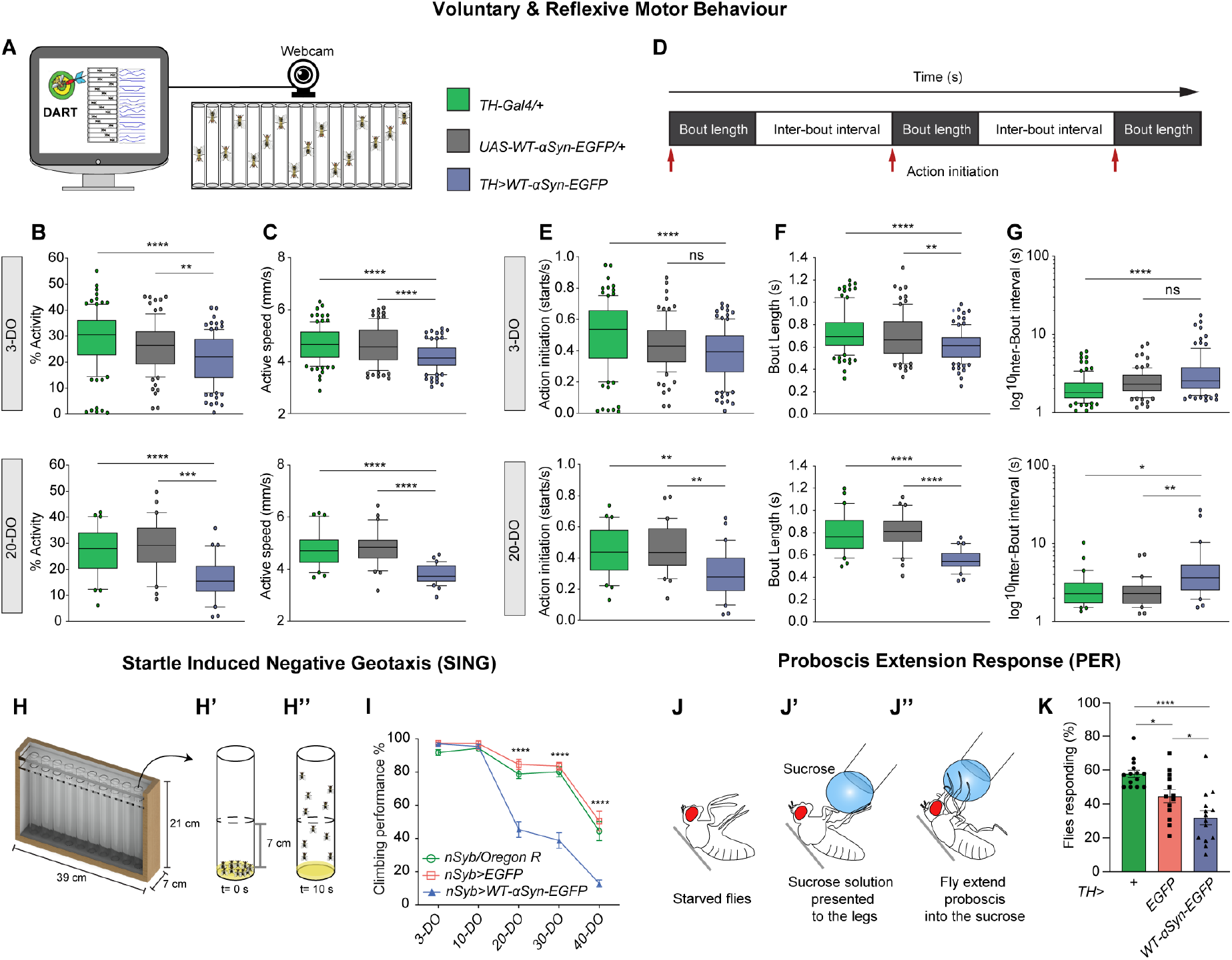
Synaptopathy induced by α-syn accumulation causes progressive motor impairment and akinetic behaviour prior to age-related dopaminergic neurodegeneration. (A) The *Drosophila* Arousal Tracking System (DART) was used to measure spontaneous activity of individual flies that were continuously recorded at 5 frames per second for 2 hours, as set by the DART software using a USB-webcam. (B) Top: spontaneous activity is reduced in 3-day-old flies (3-DO) expressing WT-α-syn-EGFP compared to controls; ****p<0.0001, **p= 0.0071 (n=90-100 flies/genotype). Bottom: spontaneous activity is further reduced in 20-day-old (20-DO) *TH*>*WT-α-syn-EGFP* when compared to controls; ****p<0.0001 and ***p= 0.0002 (n=30 flies/genotype). (C) Active speed of 3-day-old flies (3-DO) (top) and 20-day-old flies (20-DO) (top) is impaired by WT-α-syn-EGFP expression; ****p<0.0001. (D) Schematic of the activity metrics units. The initiation of a locomotor action (red arrows) is called action initiation. Bout length is the length of a motor action (dark grey boxes); and the pause between the end and beginning of a new motor action is called inter-bout interval (white boxes). (E) Top: The number of locomotor actions initiated by 3-day-old (3-DO) *TH*>*WT-α-syn-EGFP* flies was reduced compared to the control group *TH/+;* ****p<0.0001 and ns – not significant. Bottom: the locomotor actions initiated by 20-day-old (20-DO) *TH*>*WT-α-syn-EGFP* flies was reduced when compared to both controls; **p<0.0011 compared to TH/+ and **p= 0.0097 compared to *WT-α-syn-EGFP*/+ control. (F) The length of each bout of activity was shorter in *TH*>*WT-α-syn-EGFP* flies at 3 (top) and 20 (bottom) days of age, depicting their impaired ability of sustaining a locomotor action; **p= 0.0053; ****p<0.0001. (G) The length between each bout of activity was longer in flies expressing WT-α-syn-EGFP at 3 (top) and 20 (bottom) days; *p= 0.0108, **p<0.0014, ****p<0.0001. Box-and-whisker plots represent the median (horizontal line), 25% and 75% quartiles (box), and 5% and 95% quartiles (whiskers); statistical analyses were performed using Kruskal-Wallis test with Dunn’s multiple comparison post-hoc test. (H) Custom-made apparatus used to perform startle induced negative geotaxis (SING) assay which allows all fly genotypes to be probed under equal conditions simultaneously. (H’) A group of ten flies were placed in an assay tube containing 1 cm of fresh media and then allocated back in the fly holder. Next, the holder was gently tapped allowing all the flies to reach the bottom of the tubes (t=0 seconds). (H’’) After 10 seconds, the number of flies that successfully climbed above the 7cm line is recorded. (I) Cohorts of flies were analysed at 3, 10, 20, 30 and 40 days of age. The analysis showed an age-related deficiency in their climbing performance which was further enhanced by the overexpression of WT-α-syn-EGFP; ****p<0.0001; mean ± SEM are shown, n= 9-13 groups of 10 flies. (J-J’’) Proboscis extension response assay evaluated the ability of flies to respond to sucrose offer after starvation. (J) Flies were fixed in a card (grey bar) with rubber cement and were left to recover for 3 hours. (J’) Starved flies were presented with a droplet of 100 mM of sucrose to the legs and then immediately scored; (J’’) flies that extended or not their proboscis in response to sucrose were scored. (K) Young flies (5-8-day-old) expressing WT-α-syn-EGFP under control of *TH-Gal4* driver showed a reduced response to sucrose compared to controls flies, resembling an akinetic behaviour; *p= 0.03 and ****p<0.0001; mean ± SEM shown for each genotype (n= 13-14 flies/genotype).

DA-specific expression of α-syn altered activity metric parameters in 3-day-old flies (Fig. 6E-G, top and Table S1). 3-day-old *TH*>*WT-α-syn-EGFP* flies revealed a reduced ability to initiate locomotor movements compared to *TH*/+ controls (Fig. 6E, top). In addition, *TH*>*WT-α-syn-EGFP* flies showed a marked impairment in the capacity to maintain a motor action, thus showing shorter bouts of activity (Fig. 6F, top). In addition, these flies also displayed a longer interval between each activity bout (Fig. 6G, top). These alterations were also observed in 20-day-old *TH*>*WT-α-syn-EGFP* flies when compared to controls (Fig. 6E-G). Together these data demonstrate that expression of α-syn in DA neurons caused an early onset of abnormalities that affected spontaneous motor activity, exemplified by shorter bouts of activity with reduced speed and separated by longer pauses.

To investigate the impact of α-syn mediated motor impairment over an extended period of time, we utilised SING assay which probes the ability of flies to climb to the top of a tube after being gently tapped to the bottom (Miquel *et al.*, 1972; Sun *et al.*, 2018) (Fig. 6H).

Independent cohorts of flies either expressing *UAS-WT-α-syn-EGFP* or *UAS-EGFP* were aged and tested at 3, 10, 20, 30 and 40 days-old for their SING behaviour (Fig. 6I). *nSyb*>*WT-α-syn-EGFP* flies showed a severe reduction in their climbing ability starting at day 20 compared to both control groups (Fig. 6I and Table S1). This phenotype worsened as the flies reached 30 and 40 days of age, respectively (Fig. 6I and Table S1). These data demonstrate that accumulation of α-syn and in turn alterations in presynaptic proteins lead to progressive motor deficits in ageing *Drosophila*.

To further investigate the impact of α-syn expression on DA neuron function, we quantified the proboscis extension response (PER) which is modulated by a single DA neuron, the TH-VUM cell (Marella *et al.*, 2012). PER behaviour measures the proboscis extension response of flies to sugar stimuli after a short period of starvation (Fig. 6J) (Cording *et al.*, 2017). PER analysis showed the proportion of *TH*>*WT-α-syn-EGFP* flies responding to sugar stimuli was significantly lower compared to *TH*/+ and *TH>EGFP* controls (Fig. 6K and Table S1). This data demonstrates that α-syn impairs the neural response to sugar stimuli, most likely because the activity of TH-VUM neuron is diminished. Thus, indicating that impaired dopamine signalling causes akinetic behaviour in *Drosophila*.

### α-syn induced synaptopathy causes dopaminergic neurodegeneration

The synaptopathy hypothesis suggests that synaptic dysfunction and the resulting behavioural deficits precede degenerative cell loss. To test this hypothesis, we aged *TH*>*WT-α-syn-EGFP* and control flies and quantified DA neuron numbers in specific clusters of the ageing *Drosophila* brain that were identified by immunofluorescence with anti-tyrosine hydroxylase (TH) antibody (White *et al.*, 2010). We counted anti-TH-labelled DA neurons of PPL1 and PPL2, PPM1/2 and PPM3 clusters in the adult brain of 3, 20, and 40-day-old flies (Fig. 7A).

**Figure 7.**
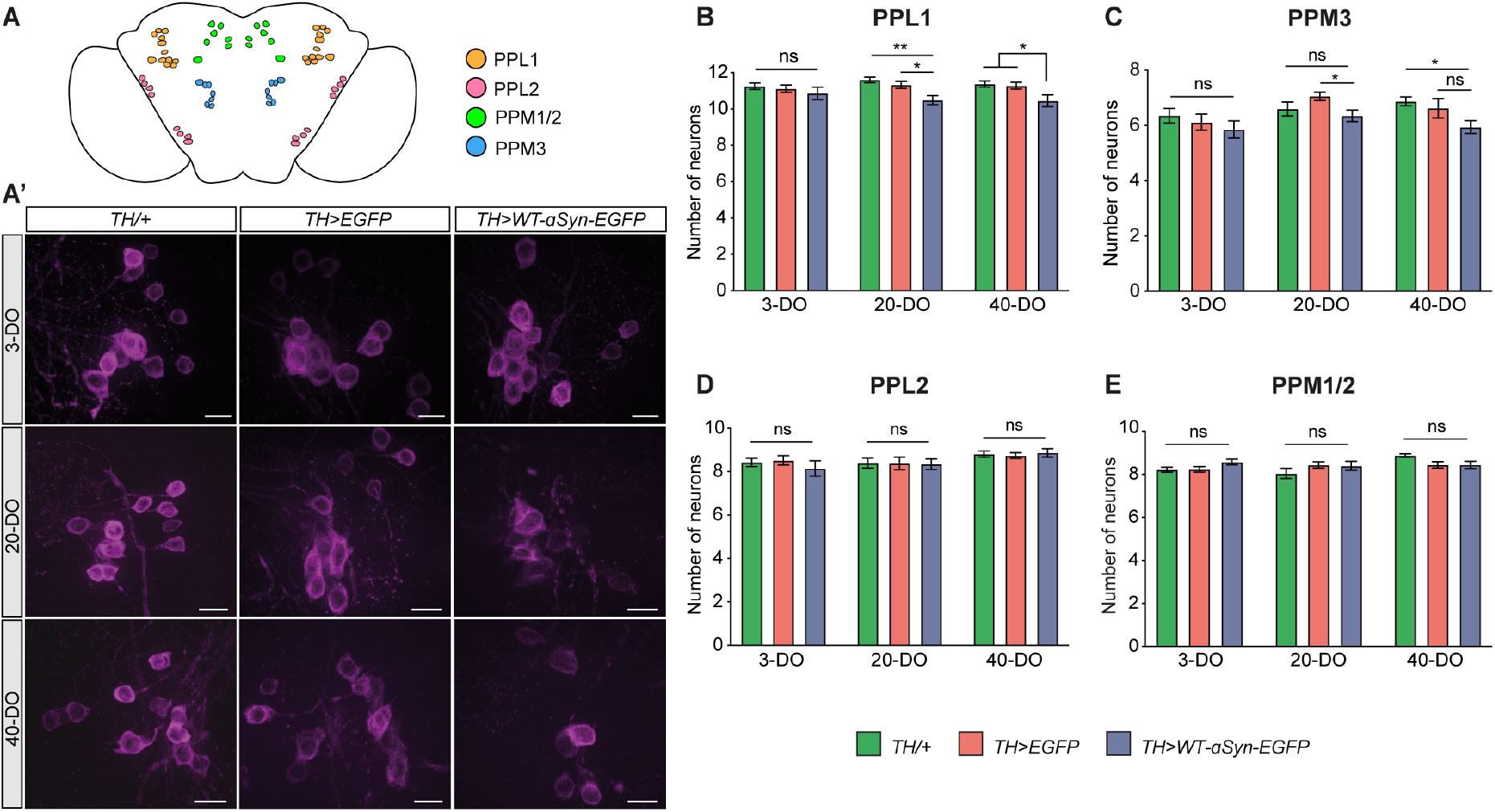
α-syn mediated synaptopathy causes progressive and age-related dopaminergic neurodegeneration. (A) Schematic depiction of dopaminergic (DA) neuron clusters in the adult *Drosophila* brain; paired posterior lateral 1 and 2 (PPL1 and PPL2), the paired posterior medial 1 and 2 (PPM1/2) and paired posterior medial 3 (PPM3). (A’) Representative iSIM images from DA neurons from PPL1 clusters immunolabelled with anti-TH antibody. (B) Number of DA neurons per hemisphere from the PPL1 cluster is reduced at day 20 and 40 compared to controls; **p= 0,0016, *p= 0,0146 for day 20; *p= 0,0207 compared to *TH*/+; *p= 0,0425 compared to *WT-α-syn-EGFP*/+ for day 40. (C) PPM3 clusters displayed a discrete loss of DA neurons in *TH*>*WT-α-syn-EGFP* flies only when compared to *TH>EGFP* at day 20 (*p=0.0292) and to *TH*/+ at day 40 (*p=0.0333). (D-E) DA neurons from PPL2 and PPM1/2 clusters were not affected by the expression of α-syn; n= 22-38 hemispheres/genotype; ns-not significant p>0.05. Mean ± SEM shown, statistical analyses were performed using one-way ANOVA with Tukey’s multiple comparison post-hoc test. Scale bar = 10 μm.

At 3 days of age, *TH*>*WT-α-syn-EGFP* flies showed no significant loss of DA neurons in the PPL1 cluster compared to the control groups (Fig. 7A’-B, Table S1). At 20-day-old, analysis of *TH*>*WT-α-syn-EGFP* flies revealed a significant loss of DA neurons compared to controls (Fig. 7A’-B, Table S1), which was also observed in 40-day-old flies compared to age-matched controls (Fig. 7A’-B, Table S1). Similar to PPL1, analysis of the PPM3 cluster revealed a discrete loss of DA neurons at day 20 when compared to *TH>EGFP* control and TH/+ control at day 40 (Fig. 7C, Table S1). In contrast to PPL1 and PPM3, PPL2 and PPM1/2 clusters showed no significant alteration in the number of DA neurons over time due to WT-α-syn overexpression (Fig. 7D-E, Table S1). Together these data demonstrate that α-syn accumulation causes region-specific and progressive degeneration of dopaminergic neurons in the ageing brain of *Drosophila*.

### Active zone protein is downregulated in human synucleinopathies

Our findings so far indicate that presynaptic α-syn accumulation caused AZ deficits that resulted in decreased neuronal function and behavioural deficits in ageing *Drosophila* that occurred prior to progressive neurodegeneration. To evaluate the clinical significance of these *in vivo* findings, we examined whether AZ proteins were also altered in patient brain. For this, we re-analysed our previously published proteomics data that compared 32 post-mortem human brains in the prefrontal cortex of patients with PDD, DLB, PD and older adults without dementia (Bereczki *et al.*, 2018). Our previous analysis identified alterations in synaptic proteins that correlated with the rate of cognitive decline and reliably discriminated PDD from Alzheimer’s disease patients (Bereczki *et al.*, 2018). Here we focused on proteins enriched in the mammalian presynaptic active zone, including homologs of the RIM, PICCOLO, ELKS and LIPRIN-α protein families (Table S2). Our initial analysis indicated potential alterations in AZ protein levels in patient brain, in particular LIPRIN-α proteins (Table S2).

Liprin-α3 and 4 proteins of the mammalian AZ play crucial roles in synapse assembly and function (Zürner and Schoch, 2009). In mammals, Liprin-α3 is highly expressed in the brain while Liprin-α4 has a lower abundance in the central nervous system (Zürner and Schoch, 2009). To corroborate our proteomics data, we carried out quantitative western blotting analysis of LIPRIN-α3 and 4 in post-mortem tissue of prefrontal cortex (BA9), anterior cingulate cortex (BA24) and parietal cortex (BA40) in control, PDD and DLB patient brain samples. Levels of LIPRIN-α3 were unaltered in all brain regions evaluated from PDD and DLB when compared to controls (Fig. 8A-C and Table S3). In contrast, expression levels of LIPRIN-α4 were significantly downregulated in the prefrontal cortex of DLB patients compared to control patients (*p=0.0349; Fig. 8B-C and Table S3). On the other hand, no significant alteration was observed in cingulate cortex (p=0.1192) or in parietal cortex (p=0.8737; Fig. 8B-C and Table S3).

**Figure 8.**
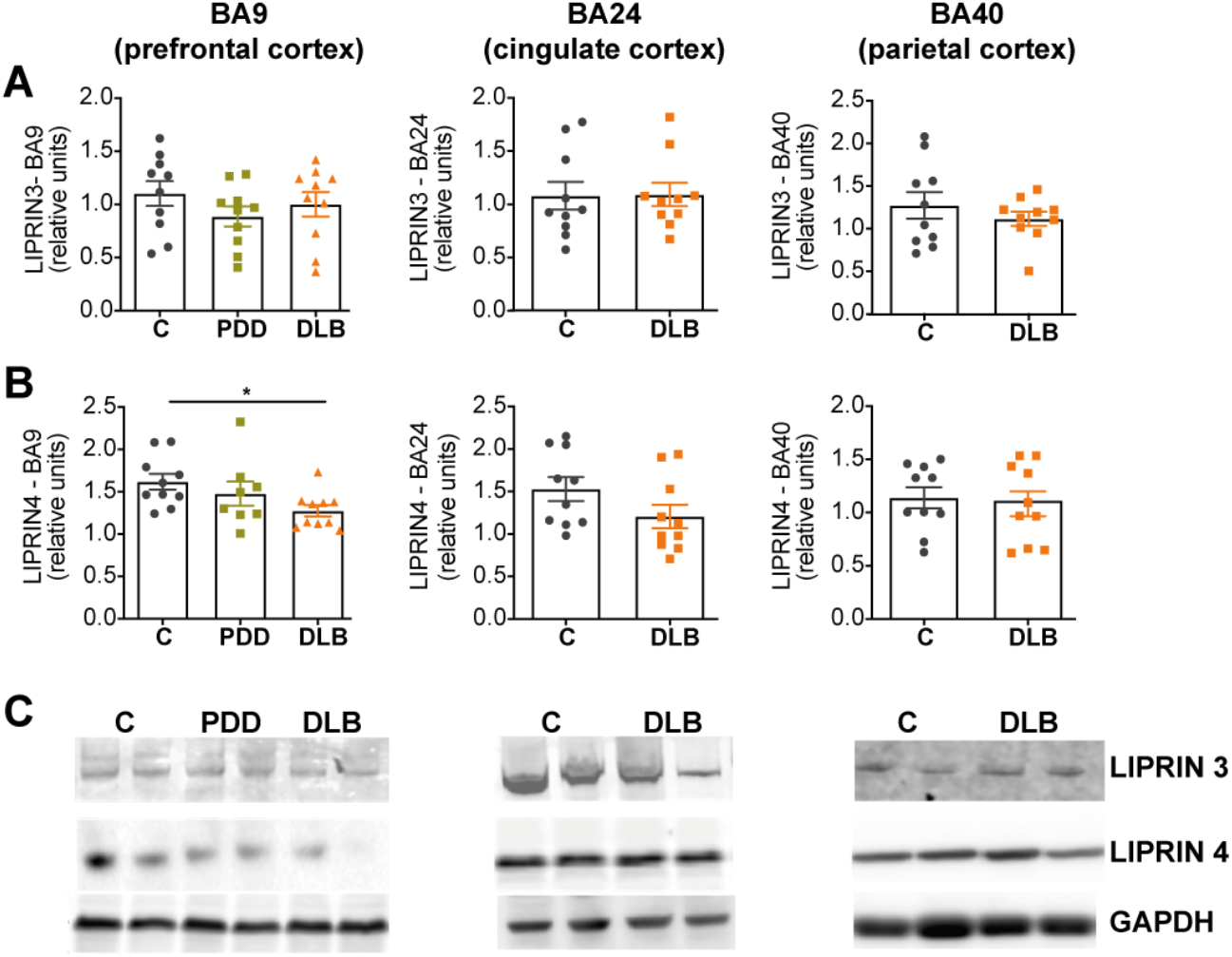
Active zone protein is affected in human synucleinopathy. (A-B) Quantitative western blots revealed no significant alteration in the expression levels of LIPRIN-α3 in the prefrontal cortex (BA9), cingulate cortex (BA24) and parietal cortex (BA40), while LIPRIN-α4 (B) was found to be downregulated in prefrontal cortex of DLB patients *p= 0.0389; (n=8-10) (C) Representative images from the western blotting membranes. All graphs are represented as mean ± SEM shown. Statistical analyses were performed using one-way ANOVA test followed by post-hoc Dunnett’s multiple comparison test.

## Discussion

α-syn accumulation and aggregation is the defining pathogenic feature of PD and other synucleinopathies (Goedert *et al.*, 2017; Bridi and Hirth, 2018). α-syn pathology correlates with synaptic deficits and subsequent synaptopathy before neurons are lost (Hornykiewicz, 1998; Calo *et al.*, 2016; Caminiti *et al.*, 2017). However, the mechanisms and cascade of events underlying the pathogenic progression from α-syn accumulation to synaptopathy and subsequent neurodegeneration are not well understood. Here we tested the hypothesis of α-syn-mediated synaptopathy by expressing the human wildtype form of α-syn in *Drosophila,* a well-established model to study PD and synucleinopathy (Feany and Bender, 2000; Auluck *et al.*, 2002), and monitored its effect over time in the ageing animal. Our findings indicate that targeted expression of human α-syn leads to its accumulation in presynaptic terminals that caused downregulation of presynaptic proteins CSP, Synapsin and Syntaxin and a reduction in the number of AZ required for synaptic transmission. In addition to synaptic alterations, α-syn accumulation caused impaired neuronal function and behavioural deficits leading to the progressive loss of dopaminergic neurons in ageing flies. These data resemble key features of the onset and progression of synucleinopathies, including PD, PDD and DLB, and demonstrate that accumulating α-syn can cause synaptopathy and progressive neurodegeneration. Our findings imply that one, if not the first cytotoxic insult of α-syn pathology, is its accumulation in presynaptic terminals, which impairs presynaptic active zones, a phenotype we also observed in post-mortem tissue of DLB patients. The resulting impaired synaptic efficacy and diminished neuronal function affect behavioural output, which over time leads to progressive neurodegeneration.

Our findings are in line with post-mortem studies where α-syn was found as small aggregates in the presynaptic terminal (Kramer and Schulz-Schaeffer, 2007; Colom-Cadena et al., 2017). Dendrites and spines of DLB patients with accumulation of aggregated α-syn were significantly smaller than those without α-syn and correlated with reduced expression levels of presynaptic proteins (Kramer and Schulz-Schaeffer, 2007). These findings in human patient material resemble synaptic phenotypes we observed as a result of targeted α-syn accumulation in *Drosophila;* they support recent experimental findings, which indicate that aggregated α-syn can directly bind and sequester presynaptic proteins (Choi *et al.*, 2018). Together with our findings, these data identify presynaptic deficits and the resultant synaptopathy as a conserved pathogenic pathway of accumulating α-syn.

In addition to alterations in presynaptic proteins, we also detected α-syn-mediated loss of BRP puncta in *Drosophila*. BRP is required for the structural integrity and function of synaptic AZs (Wagh *et al.*, 2006), responsible for vesicle docking and exocytosis of neurotransmitters (Zhai and Bellen, 2004; Wang *et al.*, 2016). Downregulation or mutational inactivation of BRP has been shown to impair T-bar formation and to reduce evoked synaptic transmission and quantal content (Wagh *et al.*, 2006). As a result, the neuronal function is affected, resulting in impaired behaviour, which is illustrated by the name *Bruchpilot*, meaning ‘crash pilot’ in German, referring to the significantly impaired manoeuvering of *BRP* mutant flies (Wagh *et al.*, 2006). Given BRPs core function, reduced numbers of BRP-positive AZs would predict reduced synaptic efficacy and impaired neural transmission in flies that accumulate α-syn in presynaptic terminals. Indeed, quantification of the SSVEP (Afsari *et al.*, 2014; Petridi *et al.*, 2020) revealed altered synaptic efficacy caused by α-syn accumulation in *TH*>*WT-α-syn-EGFP* flies. These observations in *Drosophila* are in line with a recent study in rodents which showed that overexpression of α-syn caused impairment in the electroretinogram and loss of TH positive amacrine cells in the retina (Marrocco *et al.*, 2020).

Consistent with impaired synaptic transmission, we found that adult flies expressing α-syn in DA neurons displayed behavioural deficits in spontaneous locomotor activity. 3-day-old *TH*>*WT-α-syn-EGFP* flies showed a marked reduction in activity and speed, accompanied by a significantly decreased ability to initiate and maintain motor actions, together with longer pauses between each bout of activity. These α-syn-mediated motor phenotypes became more pronounced in older flies, revealing an age-related progression of the disease that has also been observed in other *Drosophila* models of PD (Feany and Bender, 2000; Whitworth *et al.*, 2005; Humphrey *et al.*, 2012; Hindle *et al.*, 2013; Poças *et al.*, 2015; Sakai *et al.*, 2019). In addition, flies accumulating α-syn also displayed akinetic behaviour which was measured by the proboscis extension response that evaluates the response to sugar stimuli, a motor behaviour that is modulated by a single dopaminergic cell, the TH-VUM neuron (Marella *et al.*, 2012). Remarkably, *TH*>*WT-*α-syn*-EGFP* flies showed a significant reduction in their PER response, suggesting that α-syn accumulation impairs neuronal function.

This is further supported by the observed α-syn-mediated alterations in the SSVEP that is regulated by DA-rich lamina neurons in the adult brain of *Drosophila* (Afsari *et al.*, 2014; Petridi *et al.*, 2020). α-syn accumulation also caused progressive deficits in negative geotaxis, a startle induced locomotor behaviour that is controlled by dopamine in the fly brain (Feany and Bender, 2000; Riemensperger *et al.*, 2013; Poças *et al.*, 2015; Vaccaro *et al.*, 2017; Sun *et al.*, 2018; Sakai *et al.*, 2019). These findings demonstrate that α-syn-mediated synaptic alterations and impaired neurotransmission cause motor deficits in *Drosophila* affecting voluntary behaviour including action initiation and maintenance, as well as reflex activity. Comparable phenotypes have been observed in rodent models of synucleinopathies and patients with α-syn pathology (reviewed in Lashuel *et al.*, 2013; Bridi and Hirth, 2018). Together these data strongly suggest that presynaptic accumulation of α-syn causes synaptopathy and progressive behavioural deficits.

Accumulation and aggregation of α-syn are believed to cause a vicious cycle in dopaminergic neurons, triggering further accumulation of α-syn and neuronal cell death (Bridi and Hirth, 2018). Our experiments in *Drosophila* demonstrate that accumulating α-syn causes progressive degeneration of dopaminergic neurons in an age-related manner. Interestingly, α-syn accumulation preferentially affected both PPL1 and PPM3 cluster of DA neurons that regulate motor behaviour and are specifically affected in *Drosophila* models of PD (Whitworth *et al.*, 2005; Park *et al.*, 2006; Trinh *et al.*, 2008; Cunningham *et al.*, 2018). These findings resemble what is seen in PD patients where the reduction in nigrostriatal pathway connectivity occurs prior to the degenerative cell death of DA neurons in the substantia nigra pars compacta (Burke and O’Malley, 2013; Caminiti *et al.*, 2017). Of note, the progression of degenerative cell loss in *Drosophila* correlated with the progressive accumulation of α-syn levels in the ageing animal, illustrating a key characteristic of synucleinopathies, especially PD, in that specific populations of dopaminergic neurons are particularly vulnerable to α-syn burden which directly correlates with disease severity and extend of neurodegeneration (Braak *et al.*, 2002).

Alterations in synaptic proteins have been reported in clinical studies of PD, PDD and DLB patients with α-syn pathology. These studies suggest that axonal and synaptic alterations correlate with cognitive decline and the severity of the disease (Dijkstra *et al.*, 2015; Bereczki *et al.*, 2016). *In vivo* studies showed that reduced expression of α-syn was able to ameliorate neurotoxicity and behavioural deficits in conditional transgenic mice (Lim *et al.*, 2011). Furthermore, in a transgenic model of DLB/PD, pharmacological targeting of accumulating α-syn was sufficient to improve behavioural alterations and to ameliorate neurodegeneration (Wrasidlo *et al.*, 2016). More recent findings indicate that the process of Lewy Body formation, rather than fibril formation of α-syn, is linked to synaptic dysfunctions that occur before the early onset of neurodegeneration (Mahul-Mellier *et al.*, 2020). These findings are in line with our observations in *Drosophila,* which reveal insights into early pathogenesis whereby the presynaptic accumulation of α-syn affects synaptic proteins and impairs active zone-mediated neuronal function. Consistent with our findings in *Drosophila*, we found that the AZ matrix protein Liprin-α4 is downregulated in the prefrontal cortex of DLB patients which display Parkinsonian phenotypes along with dementia (Colom-Cadena *et al.*, 2017; Bereczki *et al.*, 2018). These findings suggest a pathogenic mechanism where synaptic alterations directly correlate with cognitive decline in PD, DLB and PDD patients (Bereczki *et al.*, 2018).

Taken together, our results presented here demonstrate α-syn accumulation in presynaptic terminals affects synaptic proteins and active zone integrity that impair neuronal function. The resultant synaptopathy causes behavioural deficits and progressive age-related neurodegeneration. This succession of phenotypes recapitulates key events of dying-back like neurodegeneration (Calo *et al.*, 2016; Tagliaferro and Burke, 2016; Bridi and Hirth, 2018) and provide insights into the pathogenic mechanisms underlying synaptopathy, the likely initiating event in Parkinson’s disease and related synucleinopathies.

## Materials and Methods

### Fly stocks and husbandry

All fly stocks were maintained in standard cornmeal media at 25oC in a 12 h light/dark cycle, unless for ageing experiments where flies were kept in 15% yeast/sugar media (White *et al.*, 2010; Diaper *et al.*, 2013; Solomon *et al.*, 2018). Strains used were *Oregon R, W1118, nSyb-gal4 (a kind gift from Dr Sean Sweeney), TH-gal4* (Friggi-Grelin *et al.*, 2003)*, UAS-EGFP, UAS-WT-α-syn-EGFP* (Poças *et al.*, 2015)

### Immunofluorescence

*Drosophila* larval NMJ dissections were carried out according to established protocol (Brent *et al.*, 2009) and fixed either with 3.5% formaldehyde for 25 min or Bouin’s fixative (Sigma) for 5 min. Primary antibodies used were anti-HRP (1:200 - Immunochemicals 123-605-021), anti-CSP (1:200 - DSHB), anti-Synapsin (1:50 - DSHB), anti-nSynaptobrevin (1:150 - Ohyama *et al.*, 2007; a kind gift from Dr Hugo Bellen, Baylor College of Medicine), anti-Synaptotagmin (1:1000 – West *et al.*, 2015; a kind gift from Dr Sean Sweeney, University of York), anti-SNAP-25 (1:100 - Rao *et al.*, 2001; a kind gift from Dr David Deitcher, Cornell University), anti-GFP (1:500 - Thermo Fischer A6455), anti-BRP (1:50 - DSHB). Adult CNS preparations were carried out as described previously (White *et al.*, 2010). The primary antibodies used were anti-TH (1:50 - ImmunoStar), anti-GFP (1:500 - Thermo Fischer Scientific A6455). Secondary antibodies were Alexa fluor 488 and 568 (1:150; Invitrogen); for details see the Supplementary material.

### Imaging and analysis

Z-stacks of NMJ synapses innervating muscle 6/7 of segment 3 were captured with a Nikon A1R confocal or Leica TCS SP5 microscopes. The adult *Drosophila* brain images were acquired using Nikon A1R confocal for DA neuron cluster analysis. The instant super resolution structured illumination microscopy (iSIM) was performed using Nikon Eclipse Ti-E Inverted microscope to image both for adult CNS and NMJ preparations.

For the fluorescence quantification, to build up the ratio between GFP signal in the synaptic boutons and axons, the intensity of ten synaptic boutons (labelled with anti-CSP) and ten axonal regions (positive for anti-HRP and negative for anti-CSP) were quantified per NMJ. Thus, each n number represents the average value obtained from the division of fluoresce intensity of synaptic boutons/axon in each NMJ. For fluorescence quantifications of Synapsin and CSP, z-stacks were obtained using identical settings for all genotypes with same z-axis spacing between them within the same experiment and optimised for detection without saturation of the signal (Goel *et al.*, 2017). Ten synaptic boutons were analysed per NMJ using the free hand tool from ImageJ (http://imagej.nih.gov/ij/), with each point in the graphs representing the average of ten synaptic boutons/NMJ.

BRP puncta number were manually counted in z-stacks using ImageJ and the Cell Counter plugin to record the total number of puncta per NMJ. Synapse surface area was calculated by creating a mask around the HRP channel, that labels the neuronal membrane, using ImageJ thresholding and 3D object counter (Goel *et al.*, 2017). DA neurons were manually counted through z-stacks using Cell Counter plugin using the anti-TH staining and each hemisphere represents an n number (White *et al.*, 2010).

### Western blotting

#### Drosophila heads

Quantitative Western blotting from adult fly heads were performed as previously published protocol (Solomon *et al.*, 2018). The following primary antibodies were used: anti-Synapsin (1:500 – DSHB 3C11), anti-Syntaxin (1:1000 – DSHB 8C3), anti-GFP (1:1000 - Thermo Fischer A6455), anti-beta actin (1:1000 - Abcam Ab8227), anti-beta tubulin (1:1000 – DSHB E3). Secondary antibodies were IRDye 800 conjugated goat anti-rabbit (1:10000, Rockland Immunochemicals) and Alexa Fluor 680 goat anti-mouse (1:10000, Invitrogen); for details see the Supplementary material.

### Analysis of neuronal function

The Steady State Visual Evoked Potential (SSVEP) assay measured the output of the photoreceptors and second-order lamina neurons using protocol described previously (Petridi *et al.*, 2020); for details see the Supplementary material.

### Behavioural Analyses

#### Drosophila ARousal Tracking (DART) System

DART was used to perform single fly tracking of age-matched mated females using protocol described previously (Faville *et al.*, 2015; Shaw *et al.*, 2018); for details see the Supplementary material.

#### Startle-induced negative geotaxis (SING)

SING was used to assess the locomotor ability of flies following a startle stimulus to which flies display a negative geotaxis response (modified from Ruan *et al.*, 2015). A group of ten mated age-matched female flies, per genotype, were transferred into the experimental tubes. After the tubes of all genotypes tested being placed in custom-made apparatus (see Fig. 7A), flies were allowed to acclimatise for 20 min. Control and experimental groups were always assayed together by tapping all the flies to the bottom of the tubes and allowing them to climb as a negative geotaxis response. After 10 seconds, the number of flies that successfully climbed above the 7 cm line was recorded. This assay was repeated 5 times allowing 1 min rest during between trials; for details see the Supplementary material.

#### Proboscis extension response (PER) – Akinesia assay

The PER assay was performed as protocol described previously (Petridi *et al.*, 2020); for details see the Supplementary material.

### Human post-mortem tissue analysis

#### Brain tissue samples

Detailed description of brain samples, diagnose criteria and neuropathological assessments has been previously published (Bereczki *et al.*, 2018). Brain tissue samples were provided from Brains for dementia research network. Consent for autopsy, neuropathological assessment and research were obtained and all studies were carried out under the ethical approval of the regional Ethical Review Board of Stockholm (2012/910-31/4). 30 cases in total/brain regions were used for the western blot experiments. Controls were defined as subjects with no clinical history and no neuropathological evidence of a neurodegenerative condition.

#### Quantitative Western Blotting

For western blot analysis, 500 mg of frozen tissue was homogenized in ice-cold buffer containing 50 mM Tris-HCL, 5 mM EGTA, 10 mM EDTA, protease inhibitor cocktail tablets and 2 mg/mL pepstatin A dissolved in ethanol:dimethyl sulfoxide 2:1 (Sigma). To minimize inter-blot variability, 20 μg total protein/samples were loaded in each lane of each gel on 7.5-10% SDS-polyacrylamide gel and then transferred to nitrocellulose membrane (Immobilon-P, Millipore). After blocking, membranes were incubated with primary antibodies followed by HRP conjugated secondary antibody. The following primary antibodies were used: rabbit polyclonal anti-LIPRIN-α3 (1:1000, Synaptic Systems 169 102); Rabbit anti-LIPRIN-α4 (1:1000, Abcam - ab136305); Rabbit anti-GAPDH (1:5000, Abcam, ab22555). Secondary antibodies used were: donkey anti-rabbit (1:10000, Invitrogen NA9340V) or donkey anti-rabbit (1:5000, LICOR, 926-32213); for details see the Supplementary material.

### Statistical analysis

GraphPad Prism 8 was used to perform the statistical analyses. Comparison of means were performed using either t-test; one-way ANOVA or Kruskal-Wallis test followed by Dunnett’s, Tukey’s or Dunn’s multiple comparison post-hoc tests. The significance was defined as p<0.05, error bars are shown as SEM. For complete description, please see the Supplementary material.

## Abbreviations

AZ: active zone
CSP: Cysteine String Protein
DA: dopaminergic
DART: *Drosophila* ARousal Tracking system
DLB: Dementia with Lewy bodies
iSIM: instant structured illumination microscopy
NMJ: neuromuscular junction
PD: Parkinson’s disease
PDD: Parkinson’s disease Dementia
PER: proboscis extension response
SING: startled-induced negative geotaxis
α-syn: α-synuclein

## Data availability

The data supporting the findings of this study are available within the article and in the supplementary material.

## Acknowledgements

We thank R. Faville for help with the DART system; H. Bellen, S. Sweeney and D. Deitcher for antibodies; the Bloomington stocks centre for flies; V. B. Pedrozo for the design and production of the custom-made SING apparatus; and to W. H. Au for helpful discussions and technical assistance. This work was supported by a PhD fellowship from CAPES Foundation–Ministry of Education of Brazil to JCB (BEX 13162/13-6); funding from Demensfonden, Magnus Bergwalls Stiftelse, Stohnes Stiftelse, Gamla Tjänarinnor to EB; funding from La Caixa Foundation (HR17-00595) to PMD and by the UK Medical Research Council (G0701498; MR/L010666/1) and the UK Biotechnology and Biological Sciences Research Council (BB/N001230/1) to FH.

## References

Afsari F, Christensen K V., Smith GP, Hentzer M, Nippe OM, Elliott CJH, et al. Abnormal visual gain control in a Parkinson’s disease model. Hum Mol Genet 2014; 23: 4465–4478.

Auluck PK, Chan HYE, Trojanowski JQ, Lee VMY, Bonini NM. Chaperone suppression of alpha-synuclein toxicity in a Drosophila model for Parkinson’s disease. Science 2002; 295: 865–8.

Bendor JT, Logan TP, Edwards RH. The function of α-synuclein. Neuron 2013; 79: 1044–1066.

Bereczki E, Branca RM, Francis PT, Pereira JB, Baek J-H, Hortobágyi T, et al. Synaptic markers of cognitive decline in neurodegenerative diseases: a proteomic approach. Brain 2018; 141: 582–595.

Bereczki E, Francis PT, Howlett D, Pereira JB, Höglund K, Bogstedt A, et al. Synaptic proteins predict cognitive decline in Alzheimer’s disease and Lewy body dementia. Alzheimer’s Dement 2016; 12: 1149–1158.

Braak H, Sandmann-Keil D, Gai W, Braak E. Extensive axonal Lewy neurites in Parkinson’s disease: A novel pathological feature revealed by alpha-synuclein immunocytochemistry. Neurosci Lett 1999; 265: 67–69.

Braak H, Del Tredici K, Bratzke H, Hamm-Clement J, Sandmann-Keil D, Rüb U. Staging of the intracerebral inclusion body pathology associated with idiopathic Parkinson’s disease (preclinical and clinical stages). J Neurol 2002; 249 Suppl: III/1–5.

Brent JR, Werner KM, McCabe BD. Drosophila Larval NMJ Dissection. J Vis Exp 2009: 4–5.

Bridi JC, Hirth F. Mechanisms of α-Synuclein Induced Synaptopathy in Parkinson’s Disease. Front Neurosci 2018; 12: 80.

Burgoyne RD, Morgan A. Cysteine string protein (CSP) and its role in preventing neurodegeneration. Semin Cell Dev Biol 2015; 40: 153–159.

Burke RE, O’Malley K. Axon degeneration in Parkinson’s disease. Exp Neurol 2013; 246: 72–83.

Burré J, Sharma M, Tsetsenis T, Buchman V, Etherton MR, Südhof TC. Alpha-synuclein promotes SNARE-complex assembly in vivo and in vitro. Science 2010; 329: 1663–7.

Calcagno B, Eyles D, van Alphen B, van Swinderen B. Transient activation of dopaminergic neurons during development modulates visual responsiveness, locomotion and brain activity in a dopamine ontogeny model of schizophrenia. Transl Psychiatry 2013; 3: e2026–10.

Calo L, Wegrzynowicz M, Santivañez-Perez J, Grazia Spillantini M. Synaptic failure and α-synuclein. Mov Disord 2016; 31: 169–177.

Caminiti SP, Presotto L, Baroncini D, Garibotto V, Moresco RM, Gianolli L, et al. Axonal damage and loss of connectivity in nigrostriatal and mesolimbic dopamine pathways in early Parkinson’s disease. NeuroImage Clin 2017; 14: 734–740.

Cheng HC, Ulane CM, Burke RE. Clinical progression in Parkinson disease and the neurobiology of axons. Ann Neurol 2010; 67: 715–725.

Choi M-G, Kim MJ, Kim D-G, Yu R, Jang Y-N, Oh W-J. Sequestration of synaptic proteins by alpha-synuclein aggregates leading to neurotoxicity is inhibited by small peptide. PLoS One 2018; 13: e0195339.

Colom-Cadena M, Pegueroles J, Herrmann AG, Henstridge CM, Muñoz L, Querol-Vilaseca M, et al. Synaptic phosphorylated α-synuclein in dementia with Lewy bodies. Brain 2017; 140: 3204–3214.

Cording AC, Shiaelis N, Petridi S, Middleton CA, Wilson LG, Elliott CJH. Targeted kinase inhibition relieves slowness and tremor in a Drosophila model of LRRK2 Parkinson’s disease. npj Park Dis 2017; 3: 34.

Cunningham PC, Waldeck K, Ganetzky B, Babcock DT. Neurodegeneration and locomotor dysfunction in Drosophila scarlet mutants. J Cell Sci 2018; 131: jcs216697.

Dauer W, Przedborski S. Parkinson’s disease: Mechanisms and models. Neuron 2003; 39: 889–909.

Diaper DC, Adachi Y, Lazarou L, Greenstein M, Simoes FA, Di Domenico A, et al. Drosophila TDP-43 dysfunction in glia and muscle cells cause cytological and behavioural phenotypes that characterize ALS and FTLD. Hum Mol Genet 2013; 22: 3883–93.

Dijkstra A a., Ingrassia A, de Menezes RX, van Kesteren RE, Rozemuller AJM, Heutink P, et al. Evidence for Immune Response, Axonal Dysfunction and Reduced Endocytosis in the Substantia Nigra in Early Stage Parkinson’s Disease. PLoS One 2015; 10: e0128651.

Faville R, Kottler B, Goodhill GJ, Shaw PJ, van Swinderen B. How deeply does your mutant sleep? Probing arousal to better understand sleep defects in Drosophila. Sci Rep 2015; 5: 8454.

Feany MB, Bender WW. A Drosophila model of Parkinson’s disease. Nature 2000; 404: 394–8.

Fernández-Chacón R, Wölfel M, Nishimune H, Tabares L, Schmitz F, Castellano-Muñoz M, et al. The synaptic vesicle protein CSPα prevents presynaptic degeneration. Neuron 2004; 42: 237–251.

Ferreira M, Massano J. An updated review of Parkinson’s disease genetics and clinicopathological correlations. Acta Neurol Scand 2017; 135: 273–284.

Friggi-Grelin F, Coulom H, Meller M, Gomez D, Hirsh J, Birman S. Targeted gene expression in Drosophila dopaminergic cells using regulatory sequences from tyrosine hydroxylase. J Neurobiol 2003; 54: 618–627.

Gitler D, Takagishi Y, Feng J, Ren Y, Rodriguiz RM, Wetsel WC, et al. Different presynaptic roles of synapsins at excitatory and inhibitory synapses. J Neurosci 2004; 24: 11368–80.

Goedert M, Jakes R, Spillantini MG. The Synucleinopathies: Twenty Years on. J Parkinsons Dis 2017; 7: S53–S71.

Goel P, Dickman D. Distinct homeostatic modulations stabilize reduced postsynaptic receptivity in response to presynaptic DLK signaling. Nat Commun 2018; 9: 1856.

Goel P, Li X, Dickman D. Disparate Postsynaptic Induction Mechanisms Ultimately Converge to Drive the Retrograde Enhancement of Presynaptic Efficacy. Cell Rep 2017; 21: 2339–2347.

Greengard P, Valtorta F, Czernik A, Benfenati F. Synaptic vesicle phosphoproteins and regulation of synaptic function. Science (80-) 1993; 259: 780–785.

Han J, Pluhackova K, Böckmann RA. The Multifaceted Role of SNARE Proteins in Membrane Fusion. Front Physiol 2017; 8: 5.

Hawkes CH, Del Tredici K, Braak H. A timeline for Parkinson’s disease. Park Relat Disord 2010; 16: 79–84.

Heisenberg M. Separation of receptor and lamina potentials in the electroretinogram of normal and mutant Drosophila. J Exp Biol 1971; 55: 85–100.

Hindle S, Afsari F, Stark M, Adam Middleton C, Evans GJO, Sweeney ST, et al. Dopaminergic expression of the Parkinsonian gene LRRK2-G2019S leads to non-autonomous visual neurodegeneration, accelerated by increased neural demands for energy. Hum Mol Genet 2013; 22: 2129–2140.

Hornykiewicz O. Biochemical aspects of Parkinson’s disease. Neurology 1998; 51: S2–9.

Humphrey DM, Parsons RB, Ludlow ZN, Riemensperger T, Esposito G, Verstreken P, et al. Alternative oxidase rescues mitochondria-mediated dopaminergic cell loss in Drosophila. Hum Mol Genet 2012; 21: 2698–2712.

Kalia L V, Lang AE. Parkinson’s disease. Lancet 2015; 6736: 1–17.

Kittel RJ, Wichmann C, Rasse TM, Fouquet W, Schmidt M, Schmid A, et al. Bruchpilot promotes active zone assembly, Ca2+ channel clustering, and vesicle release. Science 2006; 312: 1051–4.

Klein C, Westenberger A. Genetics of Parkinson’s disease. Cold Spring Harb Perspect Med 2012; 2: a008888.

Kouroupi G, Taoufik E, Vlachos IS, Tsioras K, Antoniou N, Papastefanaki F, et al. Defective synaptic connectivity and axonal neuropathology in a human iPSC-based model of familial Parkinson’s disease. Proc Natl Acad Sci 2017; 114: E3679–E3688.

Kramer ML, Schulz-Schaeffer WJ. Presynaptic alpha-synuclein aggregates, not Lewy bodies, cause neurodegeneration in dementia with Lewy bodies. J Neurosci 2007; 27: 1405–10.

Lang AE, Lozano AM. Parkinson’s Disease - First of two parts. N Engl J Med 1998; 339: 1044–1053.

Lang AE, Lozano AM. Parkinson’s Disease - Second of two parts. N Engl J Med 1998; 339: 1130–1143.

Lashuel H a, Overk CR, Oueslati A, Masliah E. The many faces of α-synuclein: from structure and toxicity to therapeutic target. Nat Rev Neurosci 2013; 14: 38–48.

Lim Y, Kehm VM, Lee EB, Soper JH, Li C, Trojanowski JQ, et al. α-syn suppression reverses synaptic and memory defects in a mouse model of dementia with lewy bodies. J Neurosci 2011; 31: 10076–10087.

Mahlknecht P, Seppi K, Poewe W. The concept of prodromal Parkinson’s disease. J Parkinsons Dis 2015; 5: 681–697.

Mahul-Mellier AL, Burtscher J, Maharjan N, Weerens L, Croisier M, Kuttler F, et al. The process of Lewy body formation, rather than simply α-synuclein fibrillization, is one of the major drivers of neurodegeneration. Proc Natl Acad Sci U S A 2020; 117: 4971–4982.

Marella S, Mann K, Scott K. Dopaminergic Modulation of Sucrose Acceptance Behavior in Drosophila. Neuron 2012; 73: 941–950.

Marrocco E, Indrieri A, Esposito F, Tarallo V, Carboncino A, Alvino FG, et al. α-synuclein overexpression in the retina leads to vision impairment and degeneration of dopaminergic amacrine cells. Sci Rep 2020; 10: 9619.

Mazzoni P, Shabbott B, Cortés JC. Motor control abnormalities in Parkinson’s disease. Cold Spring Harb Perspect Med 2012; 2: 1–17.

Menon KP, Carrillo RA, Zinn K. Development and plasticity of the Drosophila larval neuromuscular junction. Wiley Interdiscip Rev Dev Biol 2013; 2: 647–70.

Miquel J, Lundgren PR, Binnard R. Negative geotaxis and mating behavior in control and gamma-irradiated. Drosoph Informmation Serv 1972; 48: 48–60.

Mochizuki H, Choong CJ, Masliah E. A refined concept: α-synuclein dysregulation disease. Neurochem Int 2018; 119: 84–96.

Nippe OM, Wade AR, Elliott CJH, Chawla S. Circadian Rhythms in Visual Responsiveness in the Behaviorally Arrhythmic Drosophila Clock Mutant Clk Jrk. J Biol Rhythms 2017; 32: 583–592.

Ohyama T, Verstreken P, Ly C V., Rosenmund T, Rajan A, Tien A-C, et al. Huntingtin-interacting protein 14, a palmitoyl transferase required for exocytosis and targeting of CSP to synaptic vesicles. J Cell Biol 2007; 179: 1481–96.

Park J, Lee SB, Lee S, Kim Y, Song S, Kim S, et al. Mitochondrial dysfunction in Drosophila PINK1 mutants is complemented by parkin. Nature 2006; 441: 1157–1161.

Petridi S, Middleton CA, Ugbode C, Fellgett A, Covill L, Elliott CJH. In Vivo Visual Screen for Dopaminergic Rab ↔ LRRK2-G2019S Interactions in Drosophila Discriminates Rab10 from Rab3. Genes|Genomes|Genetics 2020; 10: 1903–1914.

Poças GM, Branco-Santos J, Herrera F, Outeiro TF, Domingos PM. Alpha-Synuclein modifies mutant huntingtin aggregation and neurotoxicity in Drosophila. Hum Mol Genet 2015; 24: 1898–1907.

Rao SS, Stewart BA, Rivlin PK, Vilinsky I, Watson BO, Lang C, et al. Two distinct effects on neurotransmission in a temperature-sensitive SNAP-25 mutant. EMBO J 2001; 20: 6761–6771.

Riemensperger T, Issa A, Pech U, Coulom H, Nguyễn M-V, Cassar M, et al. A single dopamine pathway underlies progressive locomotor deficits in a Drosophila model of Parkinson disease. Cell Rep 2013; 5: 952–60.

Ruan K, Zhu Y, Li C, Brazill JM, Zhai RG. Alternative splicing of Drosophila Nmnat functions as a switch to enhance neuroprotection under stress. Nat Commun 2015; 6: 10057.

Sakai R, Suzuki M, Ueyama M, Takeuchi T, Minakawa EN, Hayakawa H, et al. E46K mutant α-synuclein is more degradation resistant and exhibits greater toxic effects than wild-type α-synuclein in Drosophila models of Parkinson’s disease. PLoS One 2019; 14: 1–16.

Satake W, Nakabayashi Y, Mizuta I, Hirota Y, Ito C, Kubo M, et al. Genome-wide association study identifies common variants at four loci as genetic risk factors for Parkinson’s disease. Nat Genet 2009; 41: 1303–1307.

Sharma M, Burré J, Südhof TC. CSPα promotes SNARE-complex assembly by chaperoning SNAP-25 during synaptic activity. Nat Cell Biol 2011; 13: 30–9.

Shaw RE, Kottler B, Ludlow ZN, Buhl E, Kim D, Morais da Silva S, et al. In vivo expansion of functionally integrated GABAergic interneurons by targeted increase in neural progenitors. EMBO J 2018; 33: e98163.

Simón-Sánchez J, Schulte C, Bras JM, Sharma M, Gibbs JR, Berg D, et al. Genome-wide association study reveals genetic risk underlying Parkinson’s disease. Nat Genet 2009; 41: 1308–1312.

Singleton AB, Farrer M, Johnson J, Singleton A, Hague S, Kachergus J, et al. Alpha-Synuclein locus triplication causes Parkinson’s disease. Science 2003; 302: 841.

Solomon DA, Stepto A, Au WH, Adachi Y, Diaper DC, Hall R, et al. A feedback loop between dipeptide-repeat protein, TDP-43 and karyopherin-α mediates C9orf72-related neurodegeneration. Brain 2018; 141: 2908–2924.

Spillantini MG, Schmidt ML, Lee VM-Y, Trojanowski JQ, Jakes R, Goedert M. alpha-Synuclein in Lewy bodies. Nature 1997; 388: 839–840.

Südhof TC. The synaptic vesicle cycle. Annu Rev Neurosci 2004; 27: 509–547.

Südhof TC. The Presynaptic Active Zone. Neuron 2012; 75: 11–25.

Sun J, Xu AQ, Giraud J, Poppinga H, Riemensperger T, Fiala A, et al. Neural control of startle-induced locomotion by the mushroom bodies and associated neurons in drosophila. Front Syst Neurosci 2018; 12: 6.

Tagliaferro P, Burke RE. Retrograde Axonal Degeneration in Parkinson Disease. J Parkinsons Dis 2016; 6: 1–15.

Trinh K, Moore K, Wes PD, Muchowski PJ, Dey J, Andrews L, et al. Induction of the phase II detoxification pathway suppresses neuron loss in Drosophila models of Parkinson’s disease. J Neurosci 2008; 28: 465–72.

Vaccaro A, Issa A-R, Seugnet L, Birman S, Klarsfeld A. Drosophila Clock Is Required in Brain Pacemaker Neurons to Prevent Premature Locomotor Aging Independently of Its Circadian Function. PLOS Genet 2017; 13: e1006507.

Venda LL, Cragg SJ, Buchman VL, Wade-Martins R. Alpha-Synuclein and dopamine at the crossroads of Parkinson’s disease. Trends Neurosci 2010; 33: 559–568.

Volpicelli-Daley LA, Luk KC, Patel TP, Tanik SA, Riddle DM, Stieber A, et al. Exogenous α-Synuclein Fibrils Induce Lewy Body Pathology Leading to Synaptic Dysfunction and Neuron Death. Neuron 2011; 72: 57–71.

Wagh DA, Rasse TM, Asan E, Hofbauer A, Schwenkert I, Dürrbeck H, et al. Bruchpilot, a protein with homology to ELKS/CAST, is required for structural integrity and function of synaptic active zones in Drosophila. Neuron 2006; 49: 833–44.

Walker Z, Possin KL, Boeve BF, Aarsland D. Lewy body dementias. Lancet 2015; 386: 1683–1697.

Wang SSH, Held RG, Wong MY, Liu C, Karakhanyan A, Kaeser PS. Fusion Competent Synaptic Vesicles Persist upon Active Zone Disruption and Loss of Vesicle Docking. Neuron 2016; 91: 777–791.

West RJH, Lu Y, Marie B, Gao FB, Sweeney ST. Rab8, POSH, and TAK1 regulate synaptic growth in a Drosophila model of frontotemporal dementia. J Cell Biol 2015; 208: 931–947.

White KE, Humphrey DM, Hirth F. The dopaminergic system in the aging brain of Drosophila. Front Neurosci 2010; 4: 205.

Whitworth AJ, Theodore DA, Greene JC, Beneš H, Wes PD, Pallanck LJ. Increased glutathione S-transferase activity rescues dopaminergic neuron loss in a Drosophila model of Parkinson’s disease. Proc Natl Acad Sci U S A 2005; 102: 8024–8029.

Wrasidlo W, Tsigelny IF, Price DL, Dutta G, Rockenstein E, Schwarz TC, et al. A de novo compound targeting α-synuclein improves deficits in models of Parkinson’s disease. Brain 2016

Zhai RG, Bellen HJ. The architecture of the active zone in the presynaptic nerve terminal. Physiology (Bethesda) 2004; 19: 262–70.

Zinsmaier KE, Eberle KK, Buchner E, Walter N, Benzer S. Paralysis and early death in cysteine string protein mutants of Drosophila. Science 1994; 263: 977–980.

Zinsmaier KE, Hofbauer A, Heimbeck G, Pflugfelder GO, Buchner S, Buchner E. A cysteine-string protein is expressed in retina and brain of drosophila. J Neurogenet 1990; 7: 15–29.

Zürner M, Schoch S. The mouse and human Liprin-α family of scaffolding proteins: Genomic organization, expression profiling and regulation by alternative splicing. Genomics 2009; 93: 243–253.

