## Supplementary material for "Presynaptic accumulation of α-synuclein causes synaptopathy and progressive neurodegeneration"

* Correspondence to:

**This file includes:**

Figs. S1, Table S1, S2 and S3.

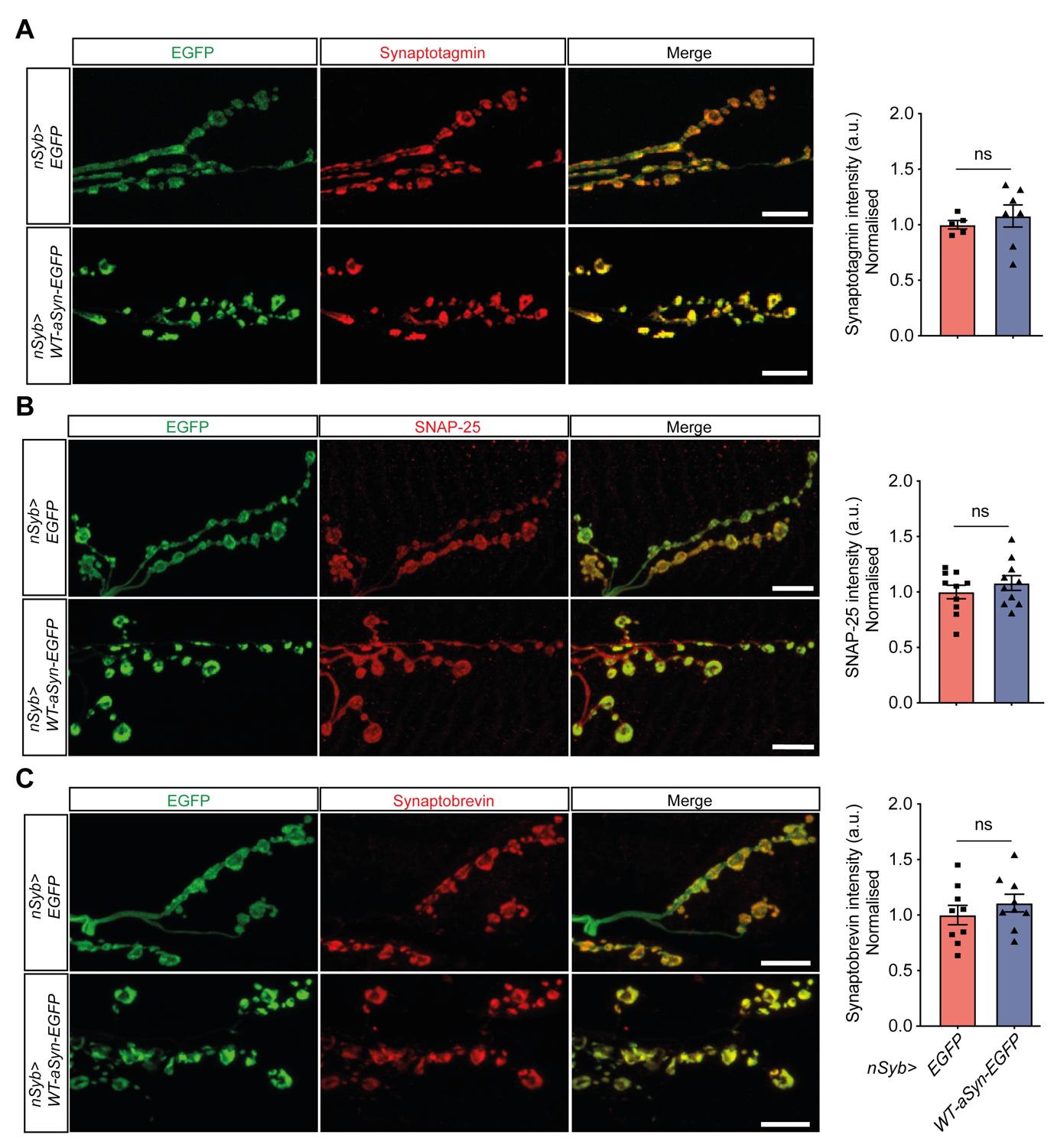

Fig. S1. Presynaptic **accumulation of α-Syn causes specific presynaptic deficits.** Representative confocal images of NMJ immunolabeled with anti-Synaptotagmin (A), anti-SNAP-25 (B), anti-nSynaptobrevin (C) and anti-GFP at the NMJ shows they are unaffected. Quantification of fluorescence intensity showed that WT-α-Syn expression under control of the pan-neuronal driver *nSyb-Gal4* caused no alterations in these proteins compared to control group expressing GFP only; ns – not significant p>0.05; n=5-10 NMJ/genotype. Mean ± SEM are shown; statistical analyses were performed using unpaired t test. Scale bars: 10 µm.

**Table S1: Quantification of phenotypes and statistical tests – Figures 1-8.**

| **Figure** | | **Genotype** | **n number** | **Mean** | **Std. Deviation** | **Std. Error of the mean** | **Statistical tests** | | | |
| --- | --- | --- | --- | --- | --- | --- | --- | --- | --- | --- |
| **Figure 1** | B | *nSyb>EGFP* | 9 | 1.152 | 0.589 | 0.197 | Unpaired two-tailed t-test | t(16)=10.40, p<0.0001 | | |
|  |  | *nSyb>WT-α-Syn-EGFP* | 9 | 23.310 | 6.362 | 2.121 |  |  |  |  |
| **Figure 2** | C | *nSyb>EGFP* | 11 | 1.000 | 0.192 | 0.058 | Unpaired two-tailed t-test | t(21)=2.332, p=0.0297 | | |
|  |  | *nSyb>WT-α-Syn-EGFP* | 12 | 0.808 | 0.202 | 0.058 |  |  |  |  |
|  | D | *nSyb>EGFP* | 21 | 1.000 | 0.316 | 0.069 |  | t(40)=11.80, p<0.0001 | | |
|  |  | *nSyb>WT-α-Syn-EGFP* | 21 | 0.166 | 0.074 | 0.016 |  |  |  |  |
| **Figure 3** | A - 3-day-old | *nSyb>EGFP* | 7 | 1.011 | 0.053 | 0.020 | Unpaired two-tailed t-test | t(14)=0.5700, p=0.5777 | | |
|  | A - 20-day-old | *nSyb>EGFP* | 9 | 1.000 | 0.024 | 0.008 |  |  |  |  |
|  | A - 3-day-old | *nSyb>WT-α-Syn-EGFP* | 8 | 1.255 | 0.066 | 0.023 |  | t(14)=2.250, p=0.0411 | | |
|  | A - 20-day-old | *nSyb>WT-α-Syn-EGFP* | 8 | 1.394 | 0.162 | 0.057 |  |  |  |  |
|  | B - 3-day-old | *nSyb>EGFP* | 3 | 1.000 | 0.003 | 0.002 |  | t(4)=3.204, p=0.0328 | | |
|  | B - 3-day-old | *nSyb>WT-α-Syn-EGFP* | 3 | 0.681 | 0.172 | 0.100 |  |  |  |  |
|  | B - 20-day-old | *nSyb>EGFP* | 6 | 1.000 | 0.098 | 0.040 |  | t(10)=4.132, p=0.002 | | |
|  | B - 20-day-old | *nSyb>WT-α-Syn-EGFP* | 6 | 0.798 | 0.068 | 0.028 |  |  |  |  |
|  | C - 3-day-old | *nSyb>EGFP* | 3 | 1.000 | 0.113 | 0.065 |  | t(4)=1.634, p=0.1776 | | |
|  | C - 3-day-old | *nSyb>WT-α-Syn-EGFP* | 3 | 0.892 | 0.017 | 0.010 |  |  |  |  |
|  | C - 20-day-old | *nSyb>EGFP* | 6 | 1.000 | 0.068 | 0.028 |  | t(10)=3.210, p=0.0093 | | |
|  | C - 20-day-old | *nSyb>WT-α-Syn-EGFP* | 6 | 0.867 | 0.075 | 0.031 |  |  |  |  |
| **Figure 4** | C | *1- nSyb/+* | 15 | 1011.0 | 136.900 | 35.350 | One-way ANOVA [F(2,43)= 4.826, p=0.0129] | Tukey's multiple comparison post-hoc test | 1 vs. 2 | Adj. p=0.994 |
|  |  | *2- nSyb>EGFP* | 15 | 1007.0 | 85.790 | 22.150 |  |  | 1 vs. 3 | Adj. p=0.0245 |
|  |  | *3- nSyb>WT-α-Syn-EGFP* | 16 | 898.0 | 118.300 | 29.580 |  |  | 2 vs. 3 | Adj. p=0.0317 |
|  | D | *1- nSyb/+* | 15 | 1.549 | 0.250 | 0.064 | One-way ANOVA [F(2,43)= 8.768, p=0.0006] |  | 1 vs. 2 | Adj. p>0.9999 |
|  |  | *2- nSyb>EGFP* | 15 | 1.549 | 0.264 | 0.068 |  |  | 1 vs. 3 | Adj. p=0.0023 |
|  |  | *3- nSyb>WT-α-Syn-EGFP* | 16 | 1.241 | 0.198 | 0.049 |  |  | 2 vs. 3 | Adj. p=0.0023 |
|  | E | *1- nSyb/+* | 15 | 130.300 | 33.390 | 8.621 | One-way ANOVA [F(2,43)= 1.643, p=0.2054] |  | 1 vs. 2 | Adj. p=0.9915 |
|  |  | *2- nSyb>EGFP* | 15 | 129.200 | 19.690 | 5.083 |  |  | 1 vs. 3 | Adj. p=0.248 |
|  |  | *3- nSyb>WT-α-Syn-EGFP* | 16 | 115.800 | 19.470 | 4.868 |  |  | 2 vs. 3 | Adj. p=0.3037 |
| **Figure 5** | E | *1- TH/+* | 9 | 8.309 | 2.559 | 0.853 | One-way ANOVA [F (2,19) = 4.859, p=0.0198] | Tukey's multiple comparison post-hoc test | 1 vs. 2 | Adj. p=0.9892 |
|  |  | *2- TH>EGFP* | 7 | 8.115 | 2.141 | 0.809 |  |  | 1 vs. 3 | Adj. p=0.0313 |
|  |  | *3-TH>WT-α-Syn-EGFP* | 6 | 12.310 | 3.553 | 1.450 |  |  | 2 vs. 3 | Adj. p=0.0325 |
|  | F | *1-TH/+* | 11 | 112.900 | 19.130 | 5.768 | One-way ANOVA [F (2,20) = 0.2507, p=0.7807] |  | 1 vs. 2 | Adj. p=0.7909 |
|  |  | *2- TH>EGFP* | 6 | 106.600 | 9.114 | 3.721 |  |  | 1 vs. 3 | Adj. p=0.9998 |
|  |  | *3- TH>WT-α-Syn-EGFP* | 6 | 113.100 | 24.970 | 10.190 |  |  | 2 vs. 3 | Adj. p=0.8261 |
| **Figure 6** | B - Top | *1- TH/+* | 100 | 28.880 | 11.030 | 1.103 | Kruskal-Wallis test | Dunn's multiple comparison post-hoc test | 1 vs. 2 | Adj. p=0.0596 |
|  |  | *2- WT-α-Syn-EGFP/+* | 89 | 25.870 | 9.572 | 1.015 |  |  | 1 vs. 3 | Adj. p<0.0001 |
|  |  | *3- TH>WT-α-Syn-EGFP* | 100 | 21.240 | 9.151 | 0.915 |  |  | 2 vs. 3 | Adj. p=0.0071 |
|  | B - Bottom | *1- TH/+* | 30 | 26.930 | 9.364 | 1.710 |  |  | 1 vs. 2 | Adj. p>0,9999 |
|  |  | *2- WT-α-Syn-EGFP/+* | 30 | 28.920 | 10.550 | 1.926 |  |  | 1 vs. 3 | Adj. p=0.0002 |
|  |  | *3- TH>WT-α-Syn-EGFP* | 30 | 16.290 | 7.953 | 1.452 |  |  | 2 vs. 3 | Adj. p<0.0001 |
|  | C - Top | *1- TH/+* | 100 | 4.657 | 0.668 | 0.067 |  |  | 1 vs. 2 | Adj. p<0,9999 |
|  |  | *2- WT-α-Syn-EGFP/+* | 89 | 4.623 | 0.725 | 0.077 |  |  | 1 vs. 3 | Adj. p<0.0001 |
|  |  | *3- TH>WT-α-Syn-EGFP* | 100 | 4.178 | 0.509 | 0.051 |  |  | 2 vs. 3 | Adj. p<0.0001 |
|  | C - Bottom | *1- TH/+* | 30 | 4.731 | 0.682 | 0.124 |  |  | 1 vs. 2 | Adj. p<0.9999 |
|  |  | *2- WT-α-Syn-EGFP/+* | 30 | 4.831 | 0.679 | 0.124 |  |  | 1 vs. 3 | Adj. p<0.0001 |
|  |  | *3- TH>WT-α-Syn-EGFP* | 30 | 3.805 | 0.387 | 0.071 |  |  | 2 vs. 3 | Adj. p<0.0001 |
|  | E- Top | *1- TH/+* | 100 | 0.509 | 0.214 | 0.021 |  |  | 1 vs. 2 | Adj. p=0.0165 |
|  |  | *2- WT-α-Syn-EGFP/+* | 89 | 0.438 | 0.162 | 0.017 |  |  | 1 vs. 3 | Adj. p<0.0001 |
|  |  | *3- TH>WT-α-Syn-EGFP* | 100 | 0.382 | 0.164 | 0.016 |  |  | 2 vs. 3 | Adj. p=0.1795 |
|  | E - Bottom | *1- TH/+* | 30 | 0.436 | 0.158 | 0.029 |  |  | 1 vs. 2 | Adj. p<0.9999 |
|  |  | *2- WT-α-Syn-EGFP/+* | 30 | 0.469 | 0.162 | 0.030 |  |  | 1 vs. 3 | Adj. p=0.0097 |
|  |  | *3- TH>WT-α-Syn-EGFP* | 30 | 0.304 | 0.156 | 0.029 |  |  | 2 vs. 3 | Adj. p=0.0011 |
|  | F - Top | *1- TH/+* | 100 | 0.734 | 0.190 | 0.019 |  |  | 1 vs. 2 | Adj. p=0.432 |
|  |  | *2- WT-α-Syn-EGFP/+* | 89 | 0.698 | 0.200 | 0.021 |  |  | 1 vs. 3 | Adj. p<0.0001 |
|  |  | *3- TH>WT-α-Syn-EGFP* | 100 | 0.608 | 0.140 | 0.014 |  |  | 2 vs. 3 | Adj. p=0.0053 |
|  | F- Bottom | *1- TH/+* | 30 | 0.801 | 0.186 | 0.034 |  |  | 1 vs. 2 | Adj. p<0.9999 |
|  |  | *2- WT-α-Syn-EGFP/+* | 30 | 0.802 | 0.162 | 0.029 |  |  | 1 vs. 3 | Adj. p<0.0001 |
|  |  | *3- TH>WT-α-Syn-EGFP* | 30 | 0.555 | 0.095 | 0.017 |  |  | 2 vs. 3 | Adj. p<0.0001 |
|  | G - Top | *1- TH/+* | 100 | 2.158 | 1.015 | 0.105 |  |  | 1 vs. 2 | Adj. p=0.0017 |
|  |  | *2- WT-α-Syn-EGFP/+* | 89 | 2.572 | 1.129 | 0.121 |  |  | 1 vs. 3 | Adj. p<0.0001 |
|  |  | *3- TH>WT-α-Syn-EGFP* | 100 | 3.504 | 2.787 | 0.280 |  |  | 2 vs. 3 | Adj. p=0.1102 |
|  | G - Bottom | *1- TH/+* | 30 | 2.871 | 1.820 | 0.332 |  |  | 1 vs. 2 | Adj. p<0.9999 |
|  |  | *2- WT-α-Syn-EGFP/+* | 30 | 2.587 | 1.438 | 0.263 |  |  | 1 vs. 3 | Adj. p=0.0108 |
|  |  | *3- TH>WT-α-Syn-EGFP* | 30 | 5.243 | 5.675 | 1.036 |  |  | 2 vs. 3 | Adj. p=0.0014 |
|  | I - 3-day-old | *1- nSyb/+* | 12 | 91.790 | 6.162 | 1.779 | One-way ANOVA [F (2, 35) =7.480, p=0.0020] | Tukey's multiple comparison post-hoc test | 1 vs. 2 | Adj. p=0.004 |
|  |  | *2- nSyb>EGFP* | 13 | 97.380 | 2.873 | 0.797 |  |  | 1 vs. 3 | Adj. p=0.0066 |
|  |  | *3- nSyb>WT-α-Syn-EGFP* | 13 | 97.080 | 2.100 | 0.583 |  |  | 2 vs. 3 | Adj. p=0.9794 |
|  | I - 10-day-old | *1- nSyb/+* | 10 | 94.400 | 4.222 | 1.335 | One-way ANOVA [F (2,27) =1.093, p=0.3496] |  | 1 vs. 2 | Adj. p=0.3239 |
|  |  | *2- nSyb>EGFP* | 10 | 97.200 | 4.638 | 1.467 |  |  | 1 vs. 3 | Adj. p=0.848 |
|  |  | *3- nSyb>WT-α-Syn-EGFP* | 10 | 95.450 | 3.947 | 1.248 |  |  | 2 vs. 3 | Adj. p=0.6359 |
|  | I - 20-day-old | *1- nSyb/+* | 10 | 78.800 | 8.651 | 2.736 | One-way ANOVA [F (2, 26) = 33.71, p<0.0001] |  | 1 vs. 2 | Adj. p=0.5097 |
|  |  | *2- nSyb>EGFP* | 9 | 84.670 | 9.434 | 3.145 |  |  | 1 vs. 3 | Adj. p<0.0001 |
|  |  | *3- nSyb>WT-α-Syn-EGFP* | 10 | 45.400 | 14.850 | 4.696 |  |  | 2 vs. 3 | Adj. p<0.0001 |
|  | I - 30-day-old | *1- nSyb/+* | 10 | 80.220 | 9.732 | 3.078 | One-way ANOVA [F (2, 27) = 49.95, p<0.0001] |  | 1 vs. 2 | Adj. p=0.7789 |
|  |  | *2- nSyb>EGFP* | 10 | 83.600 | 7.589 | 2.400 |  |  | 1 vs. 3 | Adj. p<0.0001 |
|  |  | *3- nSyb>WT-α-Syn-EGFP* | 10 | 38.800 | 14.880 | 4.706 |  |  | 2 vs. 3 | Adj. p<0.0001 |
|  | I - 40-day-old | *1- nSyb/+* | 9 | 44.440 | 17.290 | 5.762 | One-way ANOVA [F (2, 24) = 16.71, p<0.0001] |  | 1 vs. 2 | Adj. p=0.6703 |
|  |  | *2- nSyb>EGFP* | 9 | 50.440 | 17.830 | 5.942 |  |  | 1 vs. 3 | Adj. p=0.0004 |
|  |  | *3- nSyb>WT-α-Syn-EGFP* | 9 | 12.890 | 6.412 | 2.137 |  |  | 2 vs. 3 | Adj. p<0.0001 |
|  | K | *1- TH/+* | 14 | 57.830 | 8.117 | 2.169 | One-way ANOVA [F (2, 38) = 14.35, p<0.0001] |  | 1 vs. 2 | Adj. p=0.0275 |
|  |  | *2- TH>EGFP* | 13 | 44.510 | 13.870 | 3.847 |  |  | 1 vs. 3 | Adj. p<0.0001 |
|  |  | *3- TH>WT-α-Syn-EGFP* | 14 | 31.840 | 15.430 | 4.124 |  |  | 2 vs. 3 | Adj. p=0.0377 |
| **Figure 7** | B - PPL1 (3-day-old) | *1- TH/+* | 28 | 11.250 | 0.928 | 0.175 | One-way ANOVA [F (2, 79) = 0.6142, p=0.5437] | Tukey's multiple comparison post-hoc test | 1 vs. 2 | Adj. p=0.9285 |
|  |  | *2- TH>EGFP* | 26 | 11.120 | 1.033 | 0.203 |  |  | 1 vs. 3 | Adj. p=0.5223 |
|  |  | *3- TH>WT-α-Syn-EGFP* | 28 | 10.860 | 1.860 | 0.352 |  |  | 2 vs. 3 | Adj. p=0.7619 |
|  | B - PPL1 (20-day-old) | *1- TH/+* | 25 | 11.600 | 0.764 | 0.153 | One-way ANOVA [F (2, 90) = 7.474, p=0.0010] |  | 1 vs. 2 | Adj. p=0.6676 |
|  |  | *2- TH>EGFP* | 31 | 11.320 | 1.077 | 0.193 |  |  | 1 vs. 3 | Adj. p=0.0016 |
|  |  | *3- TH>WT-α-Syn-EGFP* | 37 | 10.490 | 1.502 | 0.247 |  |  | 2 vs. 3 | Adj. p=0.0146 |
|  | B - PPL1 (40-day-old) | *1- TH/+* | 32 | 11.380 | 0.942 | 0.167 | One-way ANOVA [F (2, 94) = 4,522, p=0.0133] |  | 1 vs. 2 | Adj. p=0.959 |
|  |  | *2- TH>EGFP* | 32 | 11.280 | 1.054 | 0.186 |  |  | 1 vs. 3 | Adj. p=0.0207 |
|  |  | *3- TH>WT-α-Syn-EGFP* | 33 | 10.450 | 1.872 | 0.326 |  |  | 2 vs. 3 | Adj. p=0.0425 |
|  | C - PPM3 (3-day-old) | *1- TH/+* | 28 | 6.357 | 1.393 | 0.263 | One-way ANOVA [F (2, 78) = 0.7740, p=0.4647] |  | 1 vs. 2 | Adj. p=0.8349 |
|  |  | *2- TH>EGFP* | 25 | 6.120 | 1.453 | 0.291 |  |  | 1 vs. 3 | Adj. p=0.4313 |
|  |  | *3- TH>WT-α-Syn-EGFP* | 28 | 5.857 | 1.649 | 0.312 |  |  | 2 vs. 3 | Adj. p=0.8013 |
|  | C - PPM3 (20-day-old) | *1- TH/+* | 25 | 6.600 | 1.258 | 0.252 | One-way ANOVA [F (2, 93) = 3,429, p=0.0366] |  | 1 vs. 2 | Adj. p=0.2869 |
|  |  | *2- TH>EGFP* | 34 | 7.059 | 0.886 | 0.152 |  |  | 1 vs. 3 | Adj. p=0.6808 |
|  |  | *3- TH>WT-α-Syn-EGFP* | 37 | 6.351 | 1.274 | 0.209 |  |  | 2 vs. 3 | Adj. p=0.0292 |
|  | C - PPM3 (40-day-old) | *1- TH/+* | 32 | 6.875 | 0.907 | 0.160 | One-way ANOVA [F (2, 95) = 3.497, p=0.0342] |  | 1 vs. 2 | Adj. p=0.7808 |
|  |  | *2- TH>EGFP* | 32 | 6.625 | 1.996 | 0.353 |  |  | 1 vs. 3 | Adj. p=0.0333 |
|  |  | *3- TH>WT-α-Syn-EGFP* | 34 | 5.941 | 1.369 | 0.235 |  |  | 2 vs. 3 | Adj. p=0.155 |
|  | D - PPL2 (3-day-old) | *1- TH/+* | 28 | 8.429 | 0.997 | 0.189 | One-way ANOVA [F (2, 75) = 0.5210, p=0.5961] |  | 1 vs. 2 | Adj. p=0.9682 |
|  |  | *2- TH>EGFP* | 23 | 8.522 | 1.039 | 0.217 |  |  | 1 vs. 3 | Adj. p=0.728 |
|  |  | *3- TH>WT-α-Syn-EGFP* | 27 | 8.148 | 1.854 | 0.357 |  |  | 2 vs. 3 | Adj. p=0.602 |
|  | D - PPL2 (20-day-old) | *1- TH/+* | 25 | 8.400 | 1.155 | 0.231 | One-way ANOVA [F (2, 91)= 0.0121, p=0.9879] |  | 1 vs. 2 | Adj. p=0.9989 |
|  |  | *2- TH>EGFP* | 34 | 8.382 | 1.706 | 0.293 |  |  | 1 vs. 3 | Adj. p=0.988 |
|  |  | *3- TH>WT-α-Syn-EGFP* | 35 | 8.343 | 1.434 | 0.242 |  |  | 2 vs. 3 | Adj. p=0.9932 |
|  | D - PPL2 (40-day-old) | *1- TH/+* | 32 | 8.813 | 0.821 | 0.145 | One-way ANOVA [F (2, 92) = 0.1727, p=0.8417] |  | 1 vs. 2 | Adj. p=0.9479 |
|  |  | *2- TH>EGFP* | 31 | 8.742 | 0.729 | 0.131 |  |  | 1 vs. 3 | Adj. p=0.9583 |
|  |  | *3- TH>WT-α-Syn-EGFP* | 32 | 8.875 | 1.100 | 0.194 |  |  | 2 vs. 3 | Adj. p=0.8271 |
|  | E - PPM1/2 (3-day-old) | *1- TH/+* | 28 | 8.214 | 0.630 | 0.119 | One-way ANOVA [F (2,79) = 2.602, p=0.0805] |  | 1 vs. 2 | Adj. p=0.9953 |
|  |  | *2- TH>EGFP* | 26 | 8.231 | 0.652 | 0.128 |  |  | 1 vs. 3 | Adj. p=0.1114 |
|  |  | *3- TH>WT-α-Syn-EGFP* | 28 | 8.571 | 0.690 | 0.130 |  |  | 2 vs. 3 | Adj. p=0.145 |
|  | E - PPM1/2 (20-day-old) | *1- TH/+* | 26 | 8.038 | 1.216 | 0.239 | One-way ANOVA [F (2, 96) = 1.001, p=0.3715] |  | 1 vs. 2 | Adj. p=0.3977 |
|  |  | *2- TH>EGFP* | 35 | 8.429 | 0.884 | 0.149 |  |  | 1 vs. 3 | Adj. p=0.4506 |
|  |  | *3- TH>WT-α-Syn-EGFP* | 38 | 8.395 | 1.326 | 0.215 |  |  | 2 vs. 3 | Adj. p=0.9915 |
|  | E - PPM1/2 (40-day-old) | *1- TH/+* | 32 | 8.875 | 0.421 | 0.074 | One-way ANOVA [F (2, 95) = 3.251, p=0.0431] |  | 1 vs. 2 | Adj. p=0.0752 |
|  |  | *2- TH>EGFP* | 32 | 8.438 | 0.840 | 0.149 |  |  | 1 vs. 3 | Adj. p=0.0728 |
|  |  | *3- TH>WT-α-Syn-EGFP* | 34 | 8.441 | 0.991 | 0.170 |  |  | 2 vs. 3 | Adj. p=0.9998 |

**Table S2: Human active zone core proteins in PDD and DLB patient’s vs Control**

| **Uniprot Accession nr** | **Protein name** | | **PDD vs C** | | **DLB vs C** | |
| --- | --- | --- | --- | --- | --- | --- |
|  |  |  | **FC** | **p value** | **FC** | **p value** |
| Q86UR5 | RIMS1 | RIM 1-4 | 0.85 | 0.216 | 1.08 | 0.790 |
| Q9UQ26 | RIMS2 |  | 0.89 | 0.131 | 1.02 | 0.9434 |
| Q9UJD0 | RIMS3 |  | 1.12 | 0.070 | 1.10 | 0.119 |
| Q9H426 | RIMS4 |  | 0.89 | 0.660 | 0.81 | 0.509 |
| Q9Y6V0 | PCLO | PICCOLO | 0.88 | 0.204 | 1.07 | 0.710 |
| Q8IUD2 | RB6l2/ERC1 | ELKS/CAST | 0.99 | 0.784 | 0.94 | 0.249 |
| Q13136 | LIPA1 | LIPRINα 1-4 | 0.99 | 0.923 | 0.94 | 0.572 |
| O75334 | LIPA2 |  | 0.96 | 0.827 | 0.92 | 0.635 |
| O75145 | LIPA3 |  | 0.85 | 0.182 | 0.84 | 0.125 |
| O75335 | LIPA4 |  | 0.83 | 0.131 | 0.73 | **0.0185** |

RIM: regulating synaptic membrane exocytosis; PCLO: piccolo presynaptic cytomatrix protein; RB6l2/ ERC1: ELKS/RAB6-interacting/CAST family member 1; LIPA: PTPRF interacting protein alpha.

**Table S3: Quantification of phenotypes and statistical tests – Figure 8.**

| **LIPRIN-α3 – Fig 8A** | | | | | | | |
| --- | --- | --- | --- | --- | --- | --- | --- |
|  | BA9 (prefrontal cortex) | | | BA24 (cingulate cortex) | | BA40 (parietal cortex) | |
|  | Control | PDD | DLB | Control | DLB | Control | DLB |
| Number of patients | 10 | 10 | 10 | 10 | 10 | 10 | 10 |
| Mean | 1.102 | 0.8868 | 1.001 | 1.081 | 1.092 | 1.273 | 1.118 |
| Std. Deviation | 0.3690 | 0.2963 | 0.3631 | 0.4141 | 0.3472 | 0.4958 | 0.2631 |
| Std. Error of the mean | 0.1167 | 0.0937 | 0.1148 | 0.1309 | 0.1098 | 0.1568 | 0.0832 |
| Statistical tests | One-way ANOVA [F (2.27) =0.9780, p=0.39] | | | Unpaired two-tailed t-test | | Unpaired two-tailed t-test | |
|  | Dunnett´s multiple comparison post-hoc test | | |  |  |  |  |
|  | Control vs. PDD | | Adj. p= 0.2909 | t(18)=0.068, p= 0.9462 | | t(18)=0.878, p= 0.3914 | |
|  | Control vs. DLB | | Adj. p= 0.7397 |  |  |  |  |
| **LIPRIN-α4 – Fig 8B** | | | | | | | |
|  | BA9 (prefrontal cortex) | | | BA24 (cingulate cortex) | | BA40 (parietal cortex) | |
|  | Control | PDD | DLB | Control | DLB | Control | DLB |
| Number of patients | 10 | 8 | 10 | 10 | 10 | 10 | 10 |
| Mean | 1.620 | 1.480 | 1.276 | 1.530 | 1.207 | 1.140 | 1.115 |
| Std. Deviation | 0.2977 | 0.4031 | 0.2116 | 0.4442 | 0.4384 | 0.3130 | 0.3687 |
| Std. Error of the mean | 0.0941 | 0.1425 | 0.0669 | 0.1405 | 0.1386 | 0.0989 | 0.1166 |
| Statistical tests | One-way ANOVA [F (2.25) =3.189, p=0.05] | | | Unpaired two-tailed t-test | | Unpaired two-tailed t-test | |
|  | Dunnett´s multiple comparison post-hoc test | | |  |  |  |  |
|  | Control vs. PDD | | Adj. p=0.5355 | t(18)=1.636, p=0.1192 | | t(18)=0.161, p=0.8737 | |
|  | Control vs. DLB | | Adj. p= 0.0349 |  |  |  |  |

**Detailed Materials and Methods**

**Fly stocks and husbandry**

All fly stocks were maintained in standard cornmeal media at 25^o^C in a 12 h light/dark cycle, unless for ageing experiments where flies were kept in 15% yeast/sugar media (White *et al.*, 2010; Diaper *et al.*, 2013; Solomon *et al.*, 2018). Strain used were *Oregon R, W^1118^, nSyb-gal4 (a kind gift from Dr Sean Sweeney), TH-gal4* (Friggi-Grelin *et al.*, 2003)*, UAS-EGFP, UAS-WT-α-syn-EGFP* (Poças *et al.*, 2015)

**Immunofluorescence**

*Drosophila* larval NMJ dissections were carried out according to established protocol (Brent *et al.*, 2009) and fixed either with 3.5% formaldehyde for 25 min or Bouin’s fixative (Sigma) for 5 min. Samples were blocked with 10% normal goat serum (Thermo Fisher Scientific) for 20 min and incubated with primary antibody. Primary antibodies used were anti-HRP (1:200 - Immunochemicals 123-605-021), anti-CSP (1:200 - DSHB), anti-Synapsin (1:50 - DSHB), anti-nSynaptobrevin (1:150 - Ohyama *et al.*, 2007; a kind gift from Dr Hugo Bellen, Baylor College of Medicine), anti-Synaptotagmin (1:1000 – West *et al.*, 2015; a kind gift from Dr Sean Sweeney, University of York), anti-SNAP-25 (1:100 - Rao *et al.*, 2001; a kind gift from Dr David Deitcher, Cornell University), anti-GFP (1:500 - Thermo Fischer A6455), anti-BRP (1:50 - DSHB). Adult CNS preparations were carried out as described previously (White *et al.*, 2010). The primary antibodies used were anti-TH (1:50 - ImmunoStar), anti-GFP (1:500 - Thermo Fischer Scientific A6455). Secondary antibodies were Alexa fluor 488 and 568 (1:150; Invitrogen).

**Imaging and analysis**

Z-stacks of NMJ synapses innervating muscle 6/7 of segment 3 were captured with a Nikon A1R confocal microscope with Nikon Plan Apochromat 60x NA 1.40 oil-immersion objective or Leica TCS SP5 equipped with HCX Plan Apochromat 63.0x NA 1.40 oil-immersion. The adult *Drosophila* brain images were acquired using Nikon Plan Apochromat 20x NA 0.75 objective for DA neuron cluster analysis. The instant super resolution structured illumination microscopy (iSIM) was performed using Nikon Eclipse Ti-E Inverted microscope with 100x 1.49 NA to image both for adult CNS and NMJ preparations.

For the fluorescence quantification, to build up the ratio between GFP signal in the synaptic boutons and axons, the intensity of ten synaptic boutons (labelled with anti-CSP) and ten axonal regions (positive for anti-HRP and negative for anti-CSP) were quantified per NMJ. Thus, each n number represents the average value obtained from the division of fluoresce intensity of synaptic boutons/axon in each NMJ. For fluorescence quantifications of Synapsin and CSP, z-stacks were obtained using identical settings for all genotypes with same z-axis spacing between them within the same experiment and optimised for detection without saturation of the signal (Goel *et al.*, 2017). Ten synaptic boutons were analysed per NMJ using the free hand tool from ImageJ (http://imagej.nih.gov/ij/), with each point in the graphs representing the average of ten synaptic boutons/NMJ.

BRP puncta number were manually counted in z-stacks using ImageJ and the Cell Counter plugin (developed by Dr Kurt De Vos, available at <https://imagej.nih.gov/ij/plugins/cell-counter.html>) to record the total number of puncta per NMJ. Synapse surface area was calculated by creating a mask around the HRP channel, that labels the neuronal membrane, using ImageJ thresholding and 3D object counter (Goel *et al.*, 2017). DA neurons were manually counted through z-stacks using Cell Counter plugin using the anti-TH staining and each hemisphere represents an n number (White *et al.*, 2010).

**Western blotting**

*Drosophila heads.* Quantitative Western blotting from adult fly heads were performed as previously published protocol (Solomon *et al.*, 2018). In short, adult fly heads were lysed in RIPA buffer (Sigma) containing cOmplete proteinase inhibitor (Roche) and phosSTOP phosphatase inhibitor (Roche). The total protein concentration was measured using the BCA kit (Thermo Fisher Scientific). 10 μg protein/lane was submitted to electrophoresis in 10% SDS-polyacrylamide gel and transferred to nitrocellulose membrane. Each gel contained a control lane of pooled brain homogenates used as an internal standard in all gels. After blocking with Odyssey blocking buffer (Li-COR Biosciences), the following primary antibodies used were: anti-Synapsin (1:500 – DSHB 3C11), anti-Syntaxin (1:1000 – DSHB 8C3), anti-GFP (1:1000 - Thermo Fischer A6455), anti-beta actin (1:1000 - Abcam Ab8227), anti-beta tubulin (1:1000 – DSHB E3). Secondary antibodies were IRDye 800 conjugated goat anti-rabbit (1:10000, Rockland Immunochemicals) and Alexa Fluor 680 goat anti-mouse (1:10000, Invitrogen). Membrane images were acquired Odyssey CLx Imaging System and quantified with ImageJ.

**Analysis of neuronal function**

The Steady State Visual Evoked Potential (SSVEP) assay measured the output of the photoreceptors and second-order lamina neurons. On the day of eclosion, flies were placed in the dark at 29°C, 3 day-old-flies were prepared for SSVEP measurements as described (Afsari *et al.*, 2014; Petridi *et al.*, 2020). The same protocol was used, except that stimuli were generated, and responses recorded by an Arduino Due system instead of a PC. Data was analysed in Matlab and R. Full code at <https://github.com/wadelab/flyCode>.

**Behavioural Analyses**

***Drosophila ARousal Tracking (DART).*** DART was used to perform single fly tracking of age-matched mated females (Faville *et al.*, 2015; Shaw *et al.*, 2018). During each experiment, a total of 80 flies including controls and experimental genotypes were recorded. Briefly, flies were quickly anesthetised on ice and individually placed into glass tubes. The flies were allowed to recover for 30 min at 25°C prior to the beginning of the experiment. The recording was continuously performed at 5 frames per second for 2 hours using a USB-webcam (Logitech). The x/y position of every fly was tracked and analysed using DART software in order to evaluate the relative speed and activity during the recording.

***Startle-induced negative geotaxis (SING)*.** SING was used to assess the locomotor ability of flies following a startle stimulus to which flies display a negative geotaxis response (modified from Ruan *et al.*, 2015). A group of ten mated age-matched female flies, per genotype, were selected by a mouth aspirator and transferred into the experimental tubes containing 1 cm of fresh ageing food at room temperature. After the tubes of all genotypes tested being placed in custom-made apparatus (see Fig. 7A), flies were allowed to acclimatise for 20 min prior the beginning of the assay. Control and experimental groups were always assayed together by tapping all the flies to the bottom of the tubes and allowing them to climb as a negative geotaxis response. After 10 seconds, the number of flies that successfully climbed above the 7 cm line was recorded. This assay was repeated and recorded, at 30 frames per second; 5 trials were performed for each cohort and the flies were allowed to rest during 1 min between trials. The averaged data were represented as percentage. A minimum of 9 cohorts, each consisting of 10 flies (= 90 flies in total) were tested per genotype.

***Proboscis extension response (PER)* *– Akinesia assay*.** The PER assay was recorded from 5-8-day-old flies. Flies were restrained as previously described (Cording et al., 2017) and starved at 25 °C for 3 hours before being offered a droplet of 100 mM sucrose three times. The proboscis extension responses were observed with a Grasshopper 3 (Point Grey) camera mounted on a Zeiss Stemi microscope at 200 frames/second. Each response was scored Yes/No and the median response for each sample used.

**Human post-mortem tissue analysis**

***Brain tissue samples****.* Detailed description of brain samples, diagnose criteria and neuropathological assessments has been previously published (Bereczki *et al.*, 2018)*.* Brain tissue samples were provided from Brains for dementia research network. Consent for autopsy, neuropathological assessment and research were obtained and all studies were carried out under the ethical approval of the regional Ethical Review Board of Stockholm (2012/910-31/4). 30 cases in total/brain regions were used for the western blot experiments. Controls were defined as subjects with no clinical history and no neuropathological evidence of a neurodegenerative condition.

***Quantitative Western Blotting****.* For western blot analysis, 500 mg of frozen tissue was homogenized in ice-cold buffer containing 50 mM Tris-HCL, 5 mM EGTA, 10 mM EDTA, protease inhibitor cocktail tablets (Roche, 1 tablet per 50 mL of buffer), and 2 mg/mL pepstatin A dissolved in ethanol:dimethyl sulfoxide 2:1 (Sigma). Protein concentration of each sample was measured by using BCA Protein Assay Kit (Thermo Fisher Scientific). To minimize inter-blot variability, 20 μg total protein/samples were loaded in each lane of each gel on 7.5-10% SDS-polyacrylamide gel for protein separation and then transferred to nitrocellulose membrane (Immobilon-P, Millipore). After blocking, membranes were incubated with primary antibodies followed by HRP conjugated secondary antibody. Each gel contained a control lane of pooled brain homogenates used as an internal standard. Bands were visualized using Chemiluminescent substrate (Millipore) in a LAS-3000 luminescent image reader (Fujifilm) or by using secondary antibodies compatible with the Odyssey imaging system (LI-COR Biosciences). Primary antibodies were used at the following concentrations: rabbit polyclonal anti-LIPRIN-α3 (1:1000, Synaptic Systems 169 102); Rabbit anti-LIPRIN-α4 (1:1000, Abcam - ab136305); Rabbit anti-GAPDH (1:5000, Abcam, ab22555). Secondary antibodies used were donkey anti rabbit (1:10000, Invitrogen NA9340V) or donkey anti rabbit (1:5000, LICOR, 926-32213). Western blot data were evaluated and quantified using Multi Gauge Image Analyzer (version 3.0) or Image studio Lite, respectively.

**Statistical analysis**

GraphPad Prism 8 was used to perform the statistical analyses. Comparison of means from 2 experimental conditions was performed using unpaired parametric two-tailed Student’s t-test. Comparison of means from multiple experimental conditions was performed using ANOVA, followed by Dunnett’s multiple comparison post-hoc test, when comparing the experimental groups to control only. Alternatively, when comparing all groups among each other, Tukey’s multiple comparison post-hoc test was used. Samples not normally distributed were assessed by Kruskal-Wallis test followed by Dunn’s multiple comparison post-hoc test. The significance was defined as p<0.05, error bars are shown as SEM.
